# Gulf of Mexico blue hole harbors high levels of novel microbial lineages

**DOI:** 10.1101/2020.10.18.342550

**Authors:** NV Patin, ZA Dietrich, A Stancil, M Quinan, JS Beckler, ER Hall, J Culter, CG Smith, M Taillefert, FJ Stewart

## Abstract

Exploration of oxygen-depleted marine environments has consistently revealed novel microbial taxa and metabolic capabilities that expand our understanding of microbial evolution and ecology. Marine blue holes are shallow karst formations characterized by low oxygen and high organic matter content. They are logistically challenging to sample, and thus our understanding of their biogeochemistry and microbial ecology is limited. We present a metagenomic characterization of Amberjack Hole on the Florida continental shelf (Gulf of Mexico). Dissolved oxygen became depleted at the hole’s rim (32 m water depth), remained low but detectable in an intermediate hypoxic zone (40-75 m), and then increased to a secondary peak before falling below detection in the bottom layer (80-110 m), concomitant with increases in nutrients, dissolved iron, and a series of sequentially more reduced sulfur species. Microbial communities in the bottom layer contained heretofore undocumented levels of the recently discovered phylum Woesearchaeota (up to 58% of the community), along with lineages in the bacterial Candidate Phyla Radiation (CPR). Thirty-one high-quality metagenome-assembled genomes (MAGs) showed extensive biochemical capabilities for sulfur and nitrogen cycling, as well as for resisting and respiring arsenic. One uncharacterized gene associated with a CPR lineage differentiated hypoxic from anoxic zone communities. Overall, microbial communities and geochemical profiles were stable across two sampling dates in the spring and fall of 2019. The blue hole habitat is a natural marine laboratory that provides opportunities for sampling taxa with under-characterized but potentially important roles in redox-stratified microbial processes.

## Introduction

Blue holes are subsurface caverns found in karst bedrock environments. They formed during climatic periods when low sea levels exposed the bedrock to weathering, and subsequently became submerged as sea levels rose (1). Marine blue holes differ from anchialine blue holes, such as those found in the Bahamas and the Yucatán peninsula, as they do not have freshwater layers and are not exposed to the atmosphere. Anchialine blue holes can be highly stratified, with anoxic and sulfidic bottom waters (2–4) and microbial communities distinct from other marine and freshwater systems (5–8). However, data on true marine blue holes are limited. A recent study on the Yongle Blue Hole, the deepest known marine blue hole with a bottom depth of 300 m, found the water column became anoxic around 100 m with increases in hydrogen sulfide, methane, and dissolved inorganic carbon below that depth (9). This and two additional studies also found that microbial communities in the Yongle Blue Hole were notably different from those of the surrounding pelagic water column, with anoxic layers in the hole dominated by taxa linked to sulfur oxidation and nitrate reduction (9–11). Further, deep blue hole waters may be resistant to mixing with waters outside the hole, especially in regions with limited seasonal variation or water mass intrusion. These observations suggest the potential for blue holes to harbor novel microbial lineages as a consequence of both unique geochemistry and environmental isolation.

Locations of 18 blue holes have been recorded in offshore waters on the west Florida shelf; however, many more may exist but remain undiscovered due to a lack of systematic survey (12) (J. Culter, Mote Marine Laboratory, unpublished data, December 7, 2017). According to anecdotal reports from recreational divers and fishers, the rims of these holes feature dense communities of corals, sponges, and other invertebrates, in contrast with the more barren sandy bottom of the surrounding shelf. Commercially and recreationally valuable fishes also congregate at the rims. The presence of high biomass has led to speculation that blue holes could represent offshore sources of groundwater-derived nutrients transported via extensions of the Floridan aquifer (13). However, unlike for some of the nearshore blue holes (12), no measurements of groundwater discharge have been made in the offshore blue holes. Elevated nutrient levels at these sites have broad relevance for coastal ecology in the Gulf of Mexico, particularly as they may fuel phytoplankton blooms (14,15). Indeed, the west coast of Florida experiences frequent and intense harmful algal blooms (HABs). While numerous nutrient sources have been identified as potential HAB triggers, the relative importance of these sources remains geographically unconstrained and our knowledge of what drives HABs is by no means complete (16–18).

Gulf of Mexico blue holes may be chemically stratified and devoid of oxygen. In one of the only biological studies of these features, Garman et al. (2011) explored Jewfish Sink, a coastal blue hole near Hudson, Florida. Oxygen concentrations fell from near saturation at the rim (~2 m water depth) to zero around 20 m water depth and remained below detection all the way to the bottom of the hole (~64 m water depth), with the anoxic layer further characterized by pronounced sulfide accumulation with depth. A clone library of 16S rRNA genes from microbial mats in the hole revealed a taxonomically rich community (338 operational taxonomic units, including 150 bacterial and 188 archaeal taxa), with sequences closely related to those from low-oxygen habitats including deep-sea sediments, salt marshes, cold seeps, and whale falls (6). These sequences represent taxa linked to a range of metabolisms, including dissimilatory sulfur, methane, and nitrogen cycling (both oxidative and reductive). This study also reported dense and chemically variable clouds of particulates within the hole; these included iron-sulfide minerals, suggesting the potential for microbial-metal interactions in the water column.

This work, alongside evidence from the South China Sea sites (9), suggests marine blue holes are biogeochemically complex features and potential hotspots for microbial diversity. Indeed, oxygen-depleted marine water columns have been rich targets for exploring novel microbial processes. Within the past five years alone, studies in these systems have shed new light on microbial processes of arsenic respiration (19), cryptic sulfur (20) and oxygen cycling (21), low-oxygen adapted nitrification (22) and denitrification (23), and anaerobic methane oxidation (24). Moreover, oxygen-depleted waters are expanding worldwide (25), making it critical to understand how oxygen concentration impacts microbially regulated nutrient and energy budgets, and how these impacts vary among sites. Most microbes in these systems are uncultivated and phylogenetically divergent from better-studied relatives. Many of these taxa remain unclassified beyond the phylum or class level and likely represent lineages uniquely adapted to low-oxygen conditions. In lieu of cultivation, metagenomic analysis enables exploration of both the taxonomy and function of microbial players in blue holes and similar low-oxygen environments.

Here, we used metagenomics to describe the microbial ecosystem in a marine blue hole on the west Florida shelf. To help interpret microbial processes, we also present a comprehensive electrochemistry-based analysis of redox chemical speciation. Amberjack (AJ) Hole lies ~50 km west of Sarasota, Florida at a water depth of 32 m. Like Jewfish Sink, AJ is conical in shape, with a narrow rim (25 m diameter) and a wider floor (approximately 100 m diameter, as determined by initial reconnaissance dives for this study). Water depth at the floor ranges from 110 m at the edge to 90 m at the center, where a debris pile of fine-grained sediment has accumulated. Our knowledge of AJ, like that of other marine blue holes, is limited and based primarily on exploration by a small number of technical divers. The hole’s shape and depth present a challenge for SCUBA, as well as for instrumentation or submersibles. During two expeditions in May and September 2019, we used a combination of technical divers and Niskin bottles to sample water column microbial communities. These collections spanned oxygenated waters at the rim, an anoxic but non-sulfidic intermediate layer, and a bottom layer rich in reduced sulfur compounds. Our results reveal a system that is highly stratified, apparently stable between timepoints, and phylogenetically diverse, with high representation by uncultivated and poorly understood taxa. The results suggest marine blue holes may serve as unique natural laboratories for exploring redox-stratified microbial ecosystems, while highlighting a need for future biogeochemical exploration of other blue holes and similar habitats.

## Results

### Sampling scheme

We sampled physical, chemical, and biological features of Amberjack Hole in May and September 2019. The May data set included one conductivity, temperature, and depth (CTD) profile with a coupled dissolved oxygen sensor, along with water column samples for chemical analyses (5 depths) and microbial community sequencing (11 depths, including 5 from within the hole). These samples were collected via a combination of diver bottle water exchange, hand-cast Niskin bottles, and automated Niskin sampling on a rosette (Table S1). The CTD profile was acquired by attaching a combined CTD and optical dissolved oxygen sensor to an autonomous lander deployed for 24 hours to measure sediment respiratory processes and fluxes (data not included in this study). The lander was positioned at 106 m water depth on the slope of the debris pile; the deepest May water sample was acquired by a diver at this depth.

In September, improved sampling design resulted in higher spatial resolution for all water column parameters. We obtained two CTD profiles and eight samples from within the hole for microbiome analysis. All September water samples were acquired by hand-cast Niskin bottles, with the deepest sample from 95 m, the presumed peak of the debris pile. In contrast to the May sampling, all water sampling in September was performed on the day of the CTD casts, ensuring that chemical and biological measurements were temporally coupled. We therefore focus primarily on September results (below), with exceptions where noted.

### Water column physical and chemical profiles

Amberjack Hole was highly stratified (Fig. 1). Salinity increased sharply by one PSU between the overlying water column (35.2) and the blue hole water mass (~30 m, 36.2). Within the hole, salinity decreased slightly with depth (maximum difference 0.4 PSU) until 75 m and then increased to ~36 PSU at the bottom. Dissolved oxygen decreased sharply upon entry into the hole, from 100% saturation at the rim (32 m) to less than 5% saturation by 40 m (Fig. 1B). Oxygen then increased gradually to a secondary maximum of 40% at 75 m, before dropping to near 0% (anoxia) below 80 m.

**Figure 1.**
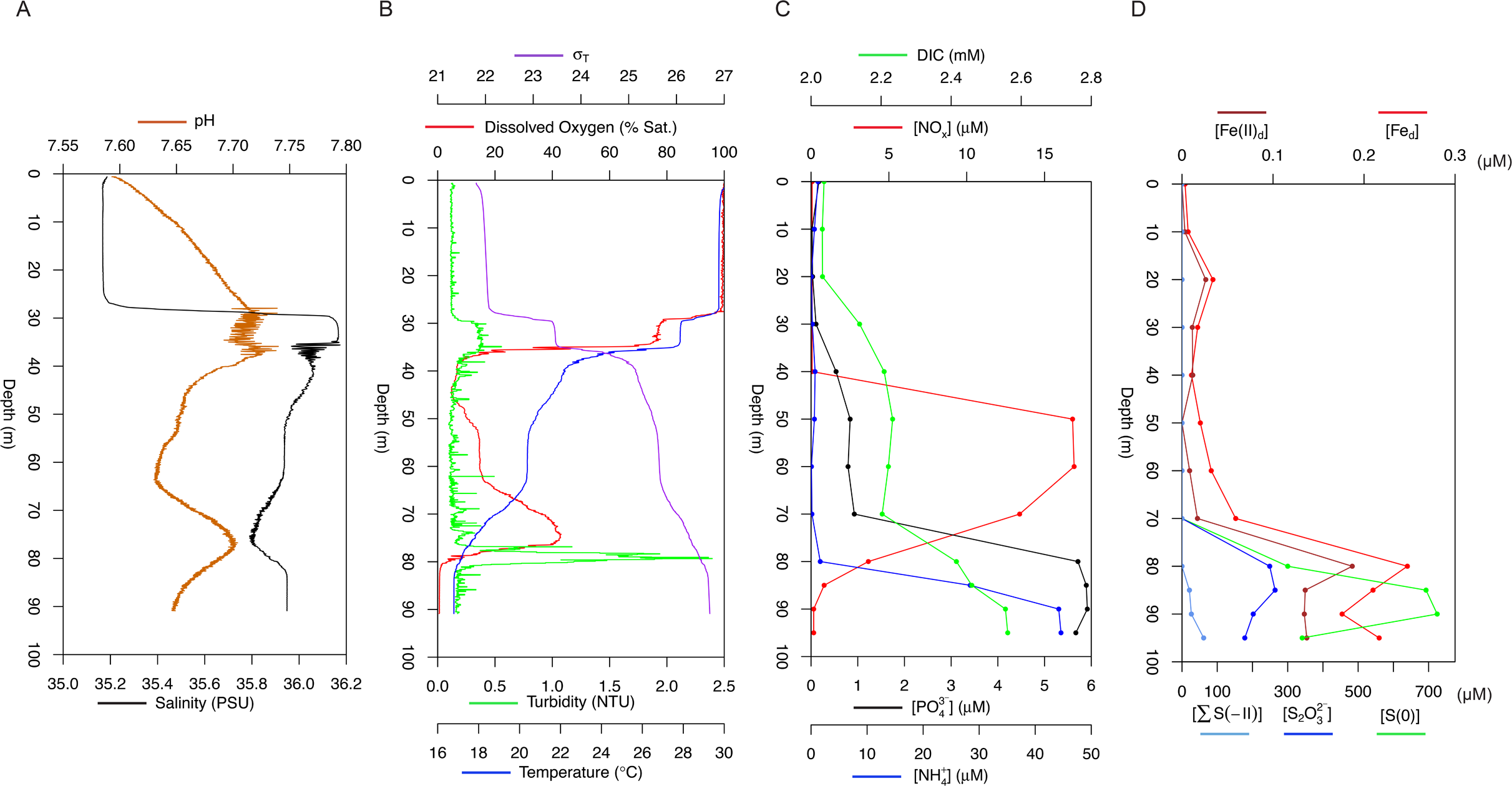
The blue hole water column in September 2019 was highly stratified. (A) Compared to the overlying water, salinity was slightly higher and pH was slightly lower inside the hole (i.e. below 30 m). A coincident dip in salinity and rise in pH was present at 75 m. (B) Dissolved oxygen concentrations varied widely, with both a primary and secondary oxycline. At 80 m, the onset of anoxia immediately below the secondary oxycline coincided with a spike in turbidity. Water density is represented by σ_T_, defined as ρ(S,T)-1000 kg m^−3^ where ρ(S,T) is the density of a sample of seawater at temperature T and salinity S, measured in kg m^−3^, at standard atmospheric pressure. (C) Dissolved inorganic carbon (DIC) increased slightly from 20 to 50 m but more intensely between 70 and 90 m, from approximately 2.2 to 2.5 mM. A sharp increase in NO_x_ (NO_2_^−^+ NO_3_^−^) between 40 and 50 m was followed by a return to near 0 between 60 and 90 m. Phosphate (PO_4_^3−^) and ammonium (NH_4_^+^) remained below 1 μm before increasing to 5-6 μm between 70 m and 80 m, respectively. (D) Dissolved ferrous iron (Fe(II)d), and total dissolved iron (Fed) increased with the transition to anoxia. Sulfur species are presented as follows: S_2_O_3_^2−^(thiosulfate, S in the +II oxidation state) peaked between 80 and 90 m. S(0) represents combined dissolved and colloidal elemental sulfur; however, the dissolved fraction may also include a small amount of polysulfide species (i.e. Sx(-II)). Finally, ∑S(-II) represents primarily hydrogen sulfide (HS^−^) but could also be minorly redundant with S(0). All iron and sulfur species increased sharply by 70-85 m, with S(0) representing the largest component of the reduced sulfur pool.

Dissolved inorganic nitrogen (nitrate (NO_3_^−^) + nitrite (NO_2_^−^), or NO_x_) spiked between 40 and 70 m from 0 to ~12 μM in May and nearly 17 μM in September, while ammonium (NH_4_^+^) and phosphate (PO_4_^3−^) increased sharply below 80 m (Fig. 1C). Bottom water nutrient concentrations were higher in September than in May, with NH_4_^+^ reaching nearly 50 μM and PO_4_^3−^ exceeding 8 μM. Particulate nutrients (N, P, and carbon (C)) and chlorophyll *a* concentrations were seemingly less variable throughout the water column in May compared to September (Fig. S2), although this pattern may be an artifact of the lower sampling resolution. In September, particulate nutrients and chlorophyll *a* spiked at the hole opening (between 30 and 40 m) with P at 0.1 μM, N at 1.2 μM, C at 11 μM, and chlorophyll *a* at nearly 1.5 μg/L. Particulate nutrients remained low deeper in the water column with the exception of P, which spiked again to 0.14 μM at 80 m. Particulate C and N also increased slightly below 80 m (Fig. S2).

Coinciding with anoxia, ferrous iron (Fe(II)d) increased from 16 nM to 187 nM between 70 and 80 m, respectively. With increasing depth, sequentially more reduced sulfur compounds were observed (Fig. 1D):

a. at 85 m, S_2_O_3_^2−^(thiosulfate, sulfur (S) in the +II oxidation state), reached a maximum of 264 μM;
b. at 90 m, S(0), representing either S_8_ (i.e. elemental sulfur, S in the 0 oxidation state) or most S within polysulfide (S_x_^2−^, e.g. S_4_^2−^ or S_8_^2−^), in filtered (< 0.7 μm) and unfiltered water reached 126 μM and 599 μM, respectively; and
c. at 95 m (deepest sample), combined ΣS(-II) (i.e. either as hydrogen sulfide (HS^−^), polysulfide (S_x_^2−^), or both) reached a maximum of 61 μM.

Due to analytical uncertainties associated with the voltammetric quantification scheme, the ∑S(-II) as presented may contribute redundantly to both (b) and (c) from a sulfur mass balance perspective. Only 6% of the S(-II) signal in the 95 m sample is lost upon filtering through 0.7 μm GFF filters (not shown), compared to 93 - 100% losses in the 85 and 80 m samples, respectively, suggesting that the deepest waters are probably enriched in true hydrogen sulfide (HS^−^). Indeed, significant HS^−^ is fluxing from sediments as revealed by benthic flux and pore water measurements (data not shown).

Notably, the zone of S(0) (i.e. elemental sulfur) accumulation between 80 and 90 m was marked by a spike in turbidity (Fig. 1B). In addition, particulate P was elevated concomitantly with Fe_d_ and Fe(II)_d_ between 70 and 80 m (Fig. 1D and S2). The latter suggests the presence of particulate or colloidal minerals and associated adsorption sites (e.g., Fe oxide colloids).

The May sampling resulted in a single data point from a depth of 85 m, which showed levels of 19.8 ± 2.3 μM S_2_O_3_^2−^, 93 μM ∑S(-II), and no S(0), similar to the September measurements from that depth.

### Microbial community composition

16S rRNA gene amplicon sequencing yielded 12,692 to 118,397 reads per sample after filtering for quality (Table S1). Water column microbial communities differed significantly based on depth grouping (PERMANOVA p=0.016), partitioning into shallow (0-32 m, oxic zone above the hole), middle (40-70 m, hypoxic zone), and deep (80-106 m, anoxic zone) groups (Fig. 2A). The shallow group was characterized by the ubiquitous cyanobacteria *Synechococcus* and *Prochlorococcus*, as well as several clades of the heterotrophic alphaproteobacterium SAR11 (particularly clade Ia) (Figure 2B). Cyanobacteria were absent below 32 m in both May and September. The middle water column featured high frequencies of *Nitrosopumilus* sp., a member of the ammonia-oxidizing Thaumarchaeota, as well as members of the sulfur-oxidizing family *Thioglobaceae* (Fig. 2B). Other groups, composing 1-10% of the community in this depth zone, included Marine Group II and III Thermoplasmatota (formerly MGII/III Euryarchaeota) and members of the family *Gimesiaceae* (phylum Planctomycetes).

**Figure 2.**
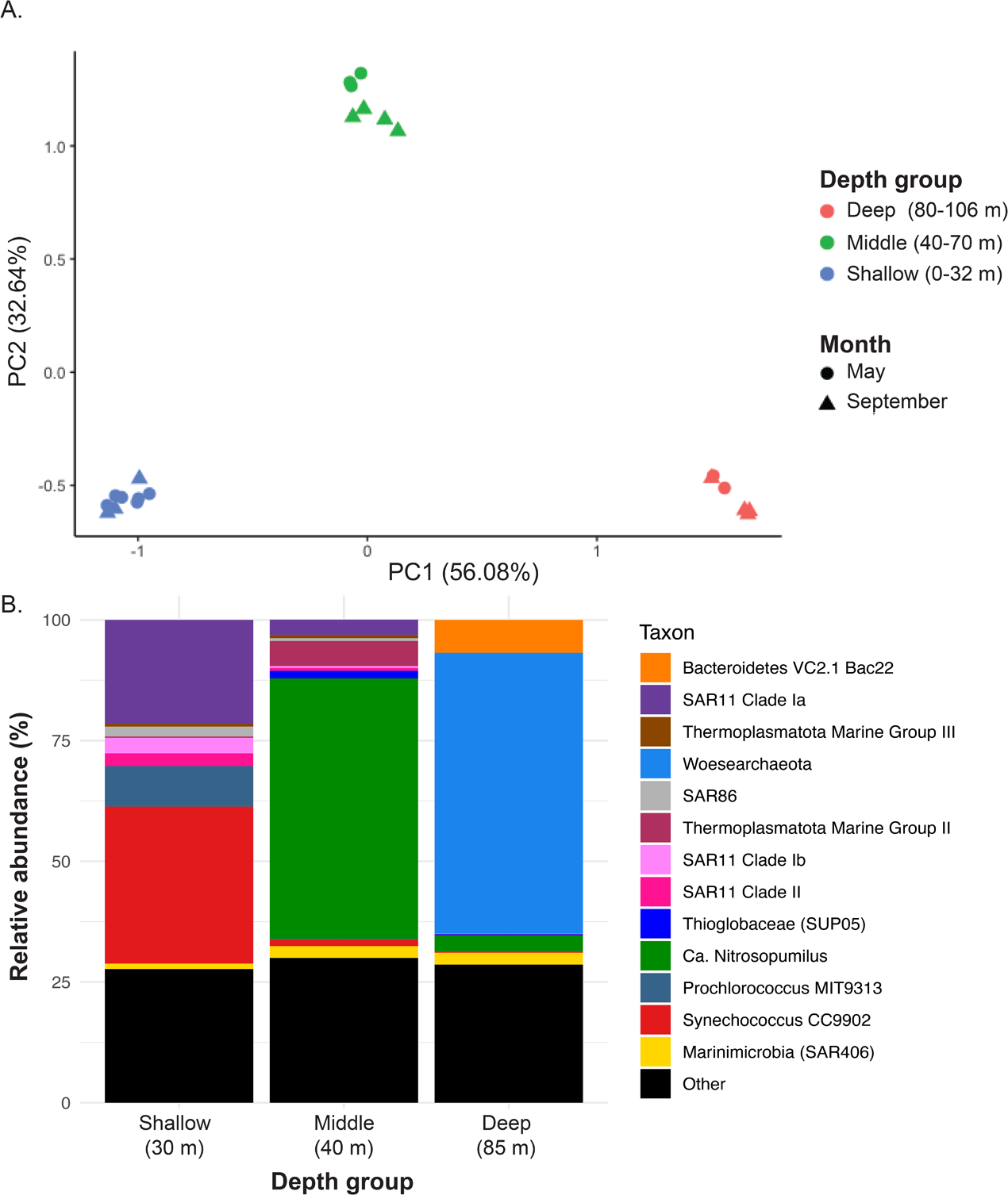
Microbial communities represented three water column depth groupings: shallow (0-32 m), middle (40-70 m), and deep (80-95 m). (A) Principal components analysis shows communities were highly similar within each depth grouping regardless of sampling date. (B) Community composition of representative samples from each of the three depth groupings show both middle and deep water column layers feature high levels (~40% frequency) of a single taxon, the ammonia-oxidizing *Nitrosopumilus* sp. in the middle water column and the Woesearchaeota in the deepest layers. Samples shown here are from September 2019 at 30 m, 40 m, and 85 m.

The anoxic and sulfidic deep water column was dominated by Woesearchaeota of the DPANN superphylum. Woesearchaeota represented 33% - 56% of all sequences below 75 m in both May and September and were comprised of 74 sequence variants (SVs), with two SVs representing between 84% (May 85 m) and 99% (September 90 m) of the total Woesearchaeotal fraction. Other well-represented groups included *Nitrosopumilus* sp, *Thioglobaceae* (SUP05 clade), and a member of the phylum Bacteroidota (formerly Bacteroidetes) associated with hydrothermal vents (“Bacteroidetes VC2.1 Bac22” in the SILVA database) (Fig. 2B). While Shannon diversity was similar among depth groups, Simpson diversity was ~50% higher in the deep compared to the middle and shallow groups (Fig. S3).

Depth patterns in community composition were highly similar between May and September. The most abundant taxa showed only minor differences in frequency between these months, with a few exceptions (Fig. 3). Marine Group II Thermoplasmatota reached 12% relative abundance in the middle hypoxic zone in September while in May this taxon never exceeded 1%; in contrast, Marine Group III peaked at around 6% in May and in September only reached about half that level. In May, *Thioglobaceae* frequency increased steadily below 30 m to peak at >10% of the community in the deep anoxic zone. In contrast, *Thioglobaceae* was not detected in the anoxic zone in September. Rather, *Arcobacter* sp. (represented by a single SV) spiked at 80 m in September to nearly 10% of the community. In all other samples, this taxon never exceeded 0.5% at any sampled depth. The 80 m depth was not sampled in May.

**Figure 3.**
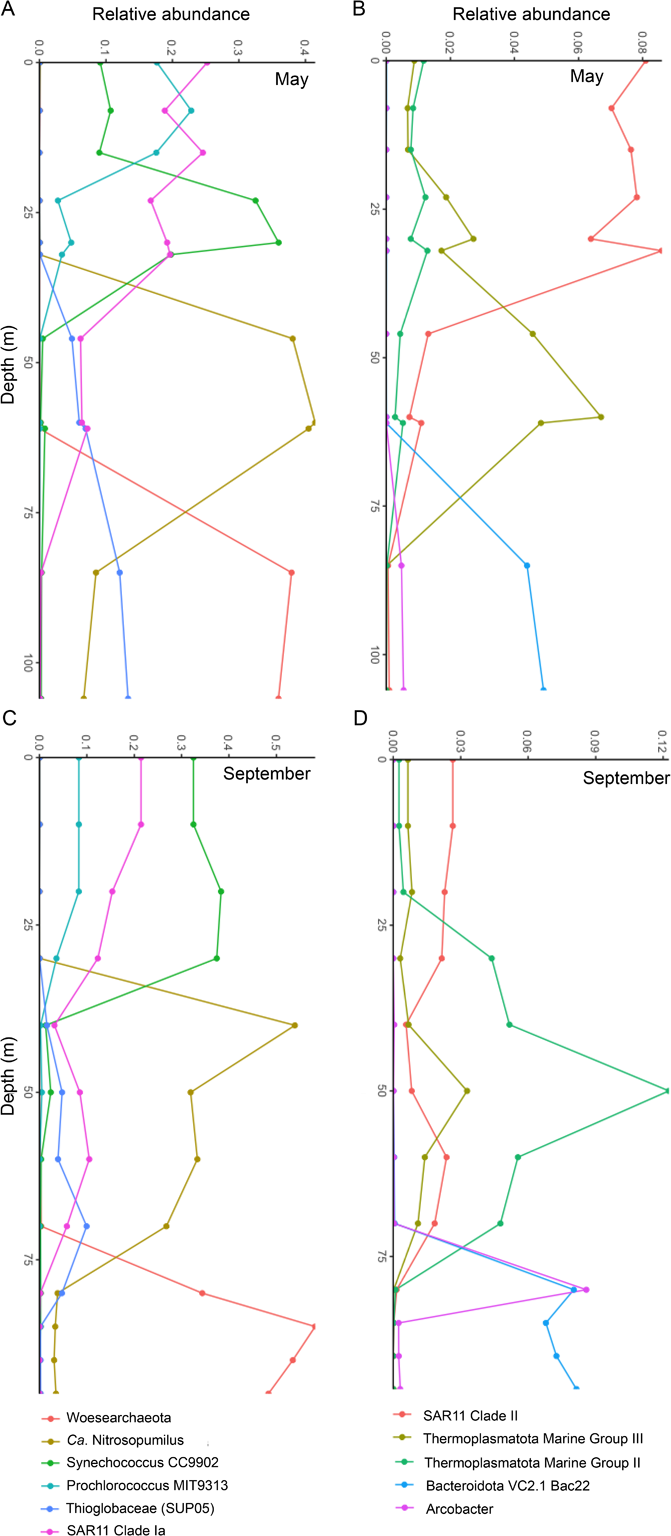
The relative abundances of the most commonly observed taxa are similar between May (A, B) and September (C, D). Notably, *Nitrosopumilus* sp. and Woesearchaeota dominated the middle and deep water column, respectively. The sulfur-oxidizing SUP05 clade increased continuously with depth in May but decreased in frequency below 70 m in September, while *Arcobacter* sp. spiked sharply at 80 m in September. Sampled depths are represented by points on each line.

### Metagenomes and MAG taxonomy

We obtained four metagenomes from two depths in both May and September. These represent communities at 60 m (both months) in the hypoxic (~15% O_2_ saturation) and *Nitrosopumilus*-dominated zone, and at 106 m (May) and 95 m (September) in the anoxic, sulfidic, and Woesearchaeota-dominated zone. Sequencing and assembly results are provided in Table S2. Metagenome binning involving data from all samples yielded 31 high-quality, non-redundant MAGs representing a diverse array of microbial taxa (Table 1). These included eight archaeal MAGs, seven of which were generated from the 60 m samples, including six belonging to the phylum Thermoplasmatota and one belonging to *Nitrosopumilus* (Thaumarchaeota). The deep metagenomes yielded two MAGs classified as members of the superphylum Patescibacteria, part of the Candidate Phyla Radiation, one of which contained a 16S rRNA gene SV classified as *Ca*. Uhrbacteria. The only archaeal MAG from the deep samples was classified as a member of the order Woesearchaeota. This MAG contained a 16S rRNA gene with 100% nucleotide identity to the most abundant SV recovered in amplicon sequencing of the bottom water communities (Table 1, Table S3). Queries against the Genome Taxonomy Database (GTDB; using MAG single-copy core genes) and SILVA database (using the 16S rRNA gene SV) classified this MAG as belonging to the order “Woesearchaeia” in the phylum Nanoarchaeota. However, this classification is outdated, as “Woesearchaeia” has recently been given the phylum-level designation Woesearchaeota (26). Only nine MAGs had a reference genome in GTDB that exceeded the minimum alignment fraction (65%); the remainder had no known close relative (Table 1).

Three MAGs representing the highest amplicon frequencies (from all recovered MAGs) were placed in phylogenies with closely related taxa, confirming their taxonomic assignments (based on GTDB and SILVA) as members of the Woesearchaeota (Fig. 4), *Nitrosopumilus* (Fig. S4), and *Thioglobaceae* (SUP05 clade) (Fig. S5). The two taxa most closely related to the Woesarchaeotal MAG are both from oxygen minimum zones in the Arabian Sea, with the next most closely related taxa from an iron-rich terrestrial hot spring and a freshwater lake (Lake Baikal). The phylogeny shows Woesearchaeotal MAGs from a wide range of environments interspersed across branches, providing weak support for clustering according to habitat type (Fig. 4). The AJ *Nitrosopumilus* MAG was most closely related to a MAG from a cold seep sponge, and relatively distantly removed from the nearest cultured *Nitrosopumilus* isolate (Fig S4). The AJ SUP05 MAG was most closely related to a clade of eight *Thioglobacaeae* spp. MAGs from hydrothermal vents, and next-most closely related to genomes associated with deep-sea invertebrates (Fig. S5).

**Figure 4.**
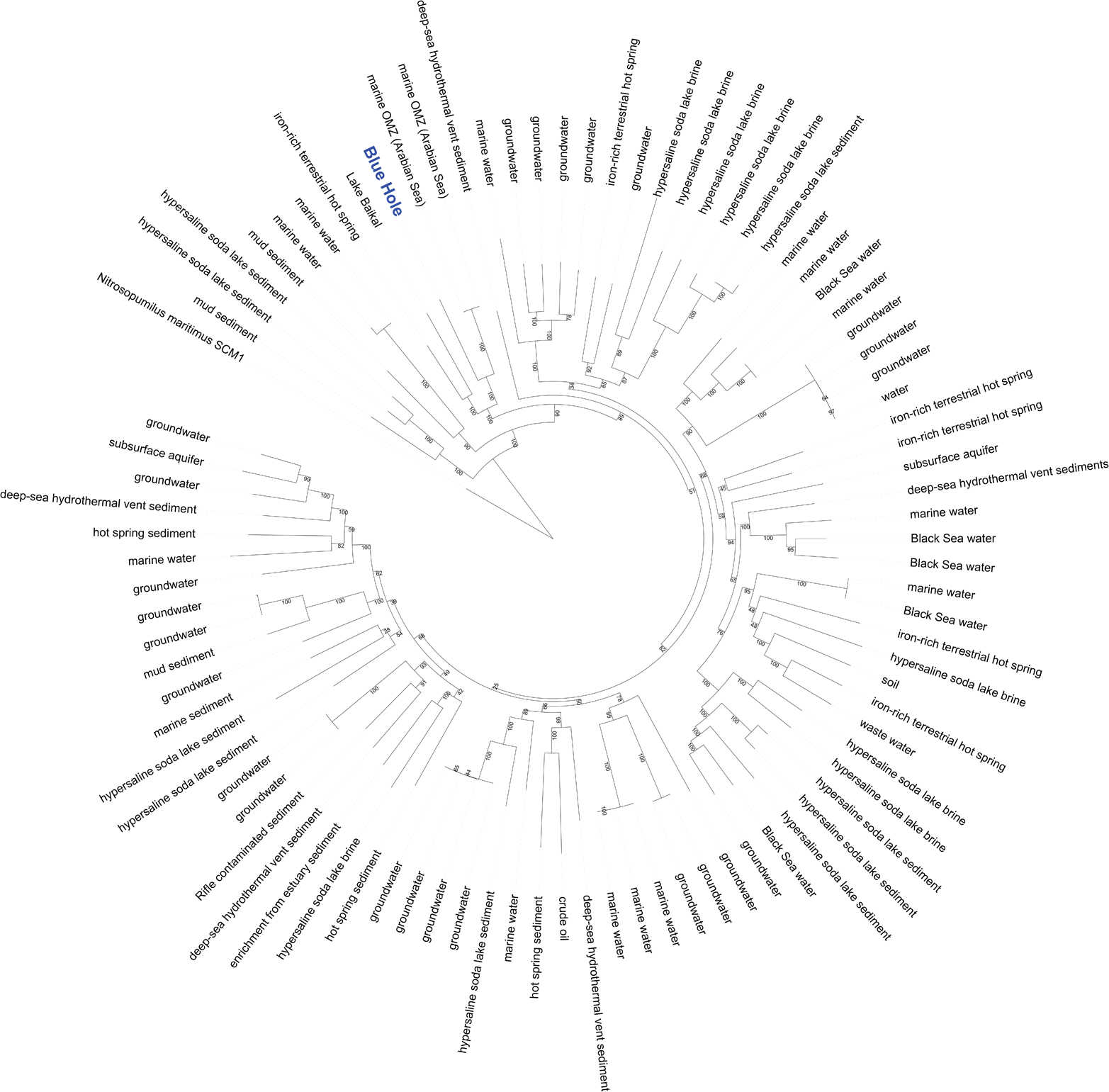
Phylogenomic analysis of the Woesearchaeotal MAG and ninety publicly available MAGs from a range of biomes show the AJ population is mostly closely related to Woesearchaeota from other marine water columns.

### Functional annotation

Based on analysis of all metagenome contigs (prior to MAG binning), broad functional categories (KEGG ‘subgroup2’ level) separated the 60 m samples from both deep samples (Fig. S6). Among the categories with a 5-fold or greater difference in frequency between the sample sets was the KEGG category ‘ECM-receptor interaction,’ which contained only two annotated genes (Fig. S7). Notably, a single gene (KO number K25373, annotated as the eukaryotic protein dentin sialophosphoprotein) occurred at much higher frequency in the deep samples relative to the 60 m samples. BLASTP queries against the NCBI nr database linked this sequence to hypothetical proteins from genomes of the candidate phylum Uhrbacteria (CPR), although these proteins shared only 30% amino acid identity with the gene in our data. This gene was identified in one of the two MAGs (BH28) classified as Patescibacteria (CPR), to which the Uhrbacteria belong. Other categories driving the sample separation included ethylbenzene degradation, bacterial chemotaxis, and o-glycan biosynthesis (Fig. S6).

Although they were not among the most differentiating categories, sulfur and nitrogen metabolism categories also differed in frequency between the depths (Fig. 5). These included genes for both assimilatory and dissimilatory metabolism. Some of the most pronounced differences in sulfur metabolism involved the dissimilatory thiosulfate (*phs*A/*psr*A) and sulfite (*dsr*AB) reductases, both of which were enriched in deep metagenomes. Sulfur metabolism genes enriched in the 60 m metagenomes also included *dmd*ABCD, which encode enzymes for the catabolism of dimethylsulfoniopropionate (DMSP).

**Figure 5.**
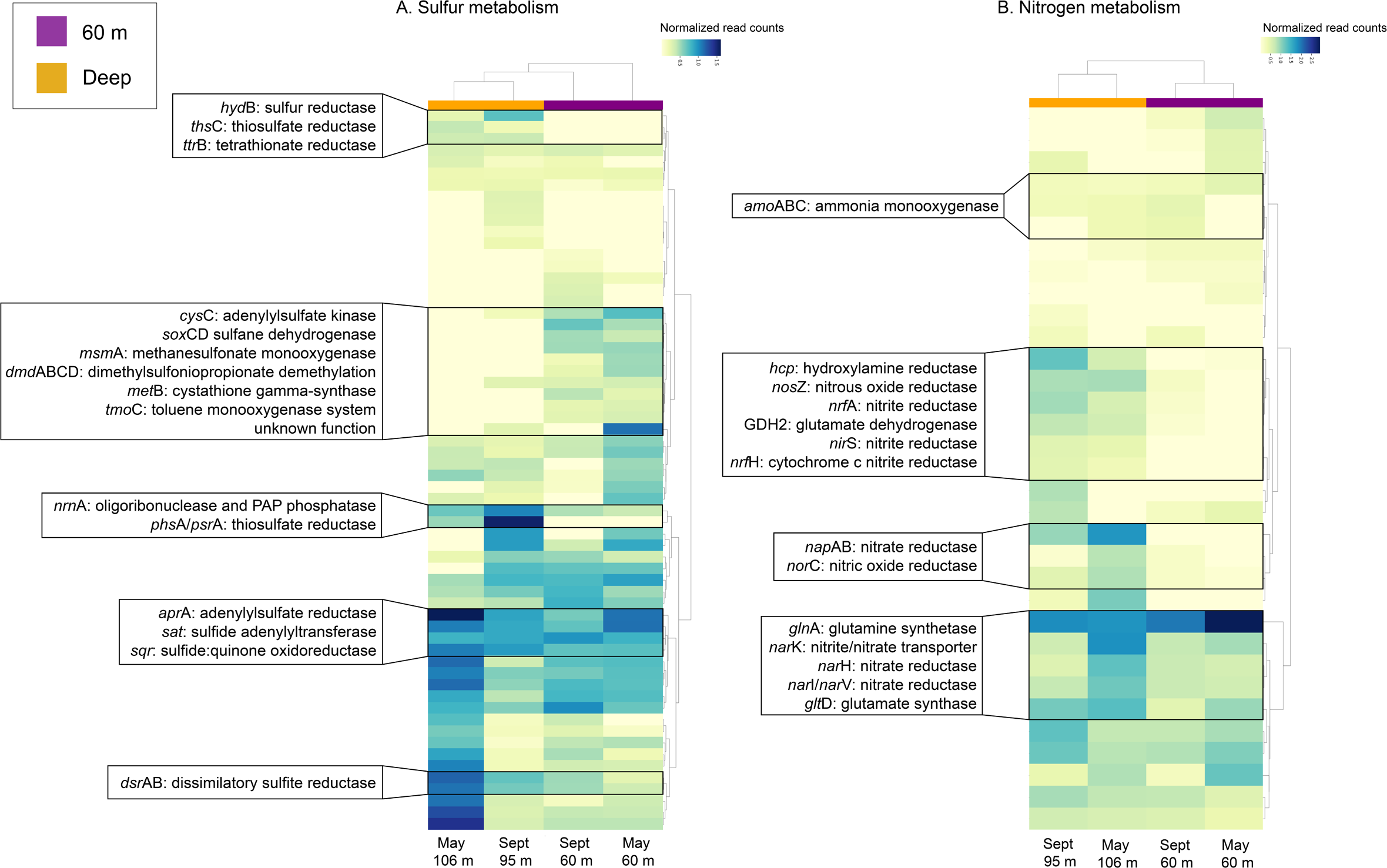
Relative abundances of genes involved in sulfur (A) and nitrogen (B) metabolism in each metagenome (May 60 m, May 106 m, September 60 m, September 95 m) show some dissimilatory genes, such as *dsr*AB and *nap*AB, are enriched in the deep water layer.

Blue hole MAGs encoded a diverse suite of metabolisms (Table 2). We detected genes involved in both reductive and oxidative pathways of dissimilatory sulfur and nitrogen cycling, arsenic respiration, methylotrophy, and carbon monoxide metabolism, among others. One MAG (BH25) is one of only two reported members of the Bacteroidota (formerly Bacteroidetes) phylum potentially capable of dissimilatory sulfur metabolism, with the other represented by a MAG from a hot spring (27). Most MAGs (23 out of 31) contained genes involved in arsenic resistance. Genes for arsenic respiration were detected in three MAGs, with two occurrences of arsenite oxidase gene *aio*A (BH16, Actinobacterial order *Microtrichales*, and BH24, alphaproteobacterial order *Rhodospirillales*) and one of arsenate reductase gene *arr*A (BH30, Desulfobacterota taxon *NaphS2*) (Fig. S8). The latter had 82% amino acid identity with a gene from another *NaphS2* strain isolated from anoxic sediment in the North Sea (DSM:14454); this gene is annotated in the FunGene database as *arr*A. Other MAG-affiliated proteins included nitrous oxide reductase (*nos*Z, in four MAGs representing three phyla), carbon monoxide dehydrogenase (*coo*FS, in BH30 (*NaphS2*, phylum Desulfobacterota)), and the ammonia and methane monooxygenases *amo*AB (in BH19, *Nitrosopumilus*) and *pmo*AB (in BH9 (family *Methylomonadaceae*, Gammaproteobacteria) (Table 2). BH9 also contained several other genes involved in C1 metabolism including *mxa*D, *mch*, *mtd*AB, and *fae* (data not shown).

The Woesearchaeotal MAG (BH21) was notable for its small size (679 Kbp, with 67% (CheckM) and 73% (anvi’o) completion values). As with other Woesearchaeotal genomes, genes for several core biosynthetic pathways were missing, including those for glycolysis/gluconeogenesis, the citric acid cycle, and the pentose phosphate pathway. Genes linked to glyoxylate and glycarboxylate metabolism and fructose and mannose metabolism were detected, along with the carbamate kinase *arc*C; however, no full energetic pathway could be reconstructed.

Only one MAG (BH24, order *Rhodospirillales*) contained genes for the full denitrification pathway (NO_3_ → N_2_). Other MAGs had the potential to perform individual steps:

NO_3_ → NO_2_, seven MAGs; NO_2_ → NO, six MAGs; NO → N_2_O, six MAGs; and N_2_O → N_2_, three MAGs (Table 2).

### Microscopy and cell counts

Microscopy-based cell counts (prokaryotes) ranged from 5 × 10^6^ to 7 × 10^6^ cells/mL within the hole (data available only from September), almost an order of magnitude lower than counts above the hole (30 m sample; Fig. S9). Counts were lowest at 80 m, the depth of the observed turbidity spike.

## Discussion

Oxygen-deficient marine water columns are crucial habitats for understanding ecosystem function under oxygen limitation and across gradients of redox substrates, representing conditions that are predicted to expand substantially in the future (25). Moreover, these systems have been a critical resource for discoveries of novel microbial diversity and unforeseen linkages between chemical cycles (19–24). While in recent years these discoveries have been facilitated by community DNA, RNA, and protein sequencing, most oxygen-depleted waters have not yet been characterized, either from an -omics perspective or via cultivation-dependent methods. This is due partly to the fact that these systems are challenging to sample and span a gradient of environmental conditions. For example, the Pacific oxygen minimum zones (OMZs) are anoxic through several hundreds of meters of the water column, cover hundreds of square kilometers of open ocean, and are relatively unaffected by processes in the underlying sediment (28). Microbial communities in these systems, especially those along the peripheries, presumably have periodic exchange with microbial communities outside the OMZ, for example via eddy intrusion, storms, or offshore transport (20). In contrast, we describe a very different oxygen-deficient zone with intense but apparently stable stratification. We show that the semi-enclosed blue hole environment exhibits a physiochemical profile and microbial community both similar to but also remarkably distinct from that of other oxygen-deficient water columns.

Amberjack Hole is characterized by an oxygen profile unlike that of most OMZs, with a secondary oxygen peak around 75 m where dissolved oxygen rose to 43% saturation before dropping back to zero (Fig. 1). In open ocean OMZs, oxygen profiles are typically unimodal, with concentrations falling along an upper oxycline, staying hypoxic or anoxic through a core layer, and then gradually increasing below the core as organic substrates are depleted with depth and microbial respiration slows (29,30). In AJ, the secondary oxygen peak at 75 m represents either a decline in net oxygen consumption driven by decreased microbial respiration, a transport-related phenomenon that affects oxygen supply, or both. Exchange of water between the offshore aquifer and blue hole water column could affect oxygen dynamics within the hole. However, the water discharging from the aquifer would be either saline connate water or saline water traveling long distances through the aquifer (13,31,32). As such, it is unlikely that oxygen would remain at levels to increase oxygen to 43% saturation. Alternatively, oxygen may come into the hole by the sinking of dense (high σ_T_, Fig 1B) oxygenated marine waters that intrude onto the shelf. Such an event could occur episodically during storms but has not been observed for this system. Turnover via severe cooling of overlying waters, or potentially upwelling and spilling of water from off the shelf, is presumably rare, and turnover by storm events unlikely. Indeed, preliminary calculations, assuming only wind-driven mixing and based on the potential energy anomaly (33), suggest that full turnover of the AJ water column would require wind speeds of 488 mph blowing for 36 hours in conjunction with 2 m/s surface current velocities in the same direction (Navid Constantinou, Australian National University, personal communication, February 2020). Thus, the observed stratification and oxygen profile may be consistent across seasons and result in a stable bottom layer that is biologically isolated from the surrounding pelagic environment.

AJ Hole is also characterized by a subsurface maximum in NO_3_^−^(nitrate) and/or NO_2_^−^(nitrite) below the main chemocline (Fig. S2), suggesting that dissolved oxygen is also consumed by nitrification. The decline in NOx coincides with the second sharp drop in oxygen concentration around 80 m. This drop is followed by the detection of reduced inorganic compounds, consistent with decreasing redox free energy expectations as a function of depth. The presence of ∑S(-II) below 80 m suggests active SO_4_^2−^ reduction. Although we detected genes potentially supporting sulfate reduction (e.g., *dsrAB*, *apr*AB), we cannot rule out the potential that the encoded proteins instead catalyze oxidative sulfur metabolism. Indeed, preliminary evidence indicates that ∑S(-II) was generated at millimolar concentrations in the sediments based on pore water profiles, and benthic lander chamber sediment flux measurements in fact show that the sediment-derived ∑S(-II) fluxes (on the order of 10 mM m^−2^ day^−1^) are comprised entirely of acid-volatile, purgeable hydrogen sulfide, ∑H2S (Beckler et al., in prep). Thus, it is likely that the reduced sulfur in the deep water column largely originates from benthic, rather than water column, sulfate reduction.

Reduced sulfur is clearly a key energy source in the deepest layers of the blue hole. We detected high concentrations of sulfur in intermediate redox states, notably S(II) (i.e. thiosulfate, S_2_O_3_^2−^) and S(0) (elemental sulfur), overlying a pronounced zone of S(-II) (hydrogen sulfide) below 80 m. Generally, oxidative sulfur metabolism proceeds with H2S/HS^−^ being oxidized to sulfite and sulfate (SO_3_^2−^/SO_4_^2−^) with S(0) and S_2_O_3_^2−^ produced as intermediates (34,35), with the relative completion depending on the pH or on the composition of the microbial community. Here, the peak concentration of S_2_O_3_^2−^ was observed at a depth above that of the peak concentration of S(0), suggesting that different steps in the overall oxidation of ∑S(-II) may be performed by various vertically-distributed niches within a complex cycle. We argue that if a single microbial population was performing complete H_2_S/HS^−^ oxidation to SO_4_^2−^(with S(0) or S(+II) as intermediates), these compounds should exhibit similar depth distributions, not stratification.

The zone of peak reduced sulfur concentration (peak concentrations of all measured sulfur compounds) occurred below 80 m (Fig. 1D) and appeared vertically decoupled from detectable dissolved oxygen, which fell below detection between 74 and 79 m (Fig. 1B). This decoupling suggests the anaerobic sulfur oxidation may also proceed with NOx as a terminal oxidant, as NO_x_ remained detectable above 90 m and decreased in concentration with depth as reduced sulfur concentrations increased (Fig. 1). Although representing only the deepest, most sulfidic layers of the blue hole, metagenomes contained diverse genes for oxidizing sequentially reduced sulfur compounds, as well as high abundances of genes for each step of the denitrification pathway (Fig. 5). Sulfur-driven denitrification is common in marine OMZs (e.g. (36)), notably in conjunction with gammaproteobacteria of the SUP05 lineage (e.g. (37)). In some cases, we detected both sulfur oxidation and denitrification genes in the same MAG, including in the SUP05 (*Thioglobaceae*) MAG (BH_2_0). We also recovered MAGs encoding only sulfur oxidation proteins or incomplete denitrification pathways (Table 2), a pattern observed in other low oxygen water columns and suggesting the likely cross-feeding of sulfur and nitrogen cycle intermediates between taxa (e.g., (38,39)). However, the zone of stratification of sulfur intermediates, roughly 70-90 m, overlaps with the zone of oxygen availability (down to 80 m) as measured with the fluorometric sensor, which has a detection limit only in the low micromolar range. Thus, it is likely that the community in this depth range consumes both oxygen and oxidized nitrogen species (and potentially other oxidants) for use in sulfur oxidation, with the relative concentrations of oxidants and sulfur intermediates potentially driving the vertical separation of microbial niches.

Interestingly, the oxic-anoxic transition at 80 m coincided with a sharp spike in turbidity. This spike was not due to an increase in microbial cell counts (Fig. S9). However, at 80 m, we recorded a sharp increase (to 10% of total) in the frequency of 16S rRNA genes assigned to *Arcobacter, a* sulfide-oxidizing bacterium known to produce filamentous sulfur at oxic-anoxic interfaces (40). Between 80 and 90 m, 70-83% of the combined elemental sulfur (S(0)) and polysulfide (∑S(-II)) signal was lost after 0.7 μm filtration, suggesting that this lost fraction was in fact elemental sulfur, as polysulfide is soluble (41,42). We did not sample at 80 m in May and therefore cannot confirm if the sharp turbidity and *Arcobacter* spikes are stable over time. However, the production of filamentous or other particulate forms of sulfur under microaerophilic conditions is consistent with the observed sulfide and oxygen gradients in both months (Fig. 1, Fig. S1). Alternatively, *Arcobacter* may be using NO_x_ to oxidize S(-II), as has been shown in denitrifying members of the genus (43).

Other oxidants may also play a role in AJ. Dissolved ferrous iron (Fe(II)) reached a maximum concentration near the turbidity spike at ~80 m, below the zone of peak O_2_/NO_x_ and above the zone of reduced sulfur (Fig. 1D). Further, total dissolved iron (Fe_d_), which includes dissolved organically-stabilized Fe(III) and FeS colloids, remained < 300 nM, but increased below 40 m, peaking just above the dynamic reduced sulfur zone (Fig. 1D). These patterns hint at a possible cryptic iron cycle, in which ∑H_2_S diffusing upwards is rapidly oxidized by dissolved Fe(III) (44) to form Fe(II), S(0), and ∑S(-II). Dissolved Fe(III) may then be recycled by reoxidation of Fe(II) by O_2_, and the S(0)/S(-II) simultaneously oxidized by O_2_ or dissolved Fe(III) to form thiosulfate, S_2_O_3_^2−^. Alternatively, ∑H_2_S may first react with Fe(II) to form the detected FeS colloids, which subsequently oxidize to form Fe(II), S(0), and S(-II). Metabolic pathways for iron oxidization (or reduction) are not easy to detect with metagenomic data, as iron metabolism genes play roles in other cellular processes and are therefore suggestive but not diagnostic of iron metabolism (45,46). Nonetheless, additional metagenomic sampling at finer spatial resolution may help identify a role for microbial activity in blue hole iron cycling. The oxidation of iron, however, is rapid even abiotically, and the vertical distribution of iron in this system can partially explain the apparent vertical disconnect between the ∑S(-II) and O_2_. Indeed, this phenomenon has been observed in a nearshore Florida hole (6), in which FeS colloids formed at the interface between 30 and 50 m. AJ appears to be enriched instead in colloidal S(0), with only nanomolar concentrations of FeS colloids.

The pronounced stratification of blue hole chemistry is consistent with differentiation of microbial communities with depth. In both May and September, microbiome composition varied distinctly among oxic, hypoxic, and anoxic, sulfidic layers (Fig. 2). Shannon diversity was similar among all communities, but Simpson diversity was notably higher in the anoxic depth group (Fig. S3), reflecting the lower evenness of these microbiomes. Surprisingly, the deepest communities were dominated by a recently described archaeal lineage, the Woesearchaeota (47). Woesearchaeota have been detected in a wide variety of biomes, including groundwater, terrestrial and marine sediments, wetlands, deep-sea hydrothermal vents, and hypersaline lakes (48). However, their relative abundance is consistently low, at most ~5% of the total microbial community in any given environment, with the highest proportions observed in freshwater sediments (48) and high-altitude lakes (49). In contrast, Woesearchaeota comprised at least one third of the blue hole microbiome between 75 and 106 meters, reaching a maximum of nearly 60% in the anoxic sulfidic layer in September (Fig. 2, Fig. 3). Remarkably, one 16S rRNA sequence variant represented up to 97% (and only two SVs represented up to 99%) of all Woesearchaeotal amplicons in each sample, suggesting low intrapopulation diversity. Dominance (>50% of the community) by a single strain variant is relatively uncommon in pelagic marine microbiomes; it is even more rare that such occurrences involve members of the Archaea. The AJ Woesearchaeotal MAG, which contained the dominant 16S rRNA Woesearchaeota amplicon sequence, was 679 Kbp and similar in size to other Woesearchaeotal genomes, which rarely exceed 1 Mbp (47,48). This MAG was most closely related to MAGs from an OMZ in the Arabian Sea (Fig. 5), suggesting the potential for a Woesearchaeotal clade specific to a pelagic, marine, low oxygen niche. At around 70% completion, the AJ MAG did not contain genes of metabolic processes common to marine low oxygen systems, notably dissimilatory sulfur or nitrogen cycling, anaerobic metabolism, or autotrophy. Indeed, consistent with characterizations of other Woesearchaeotal genomes (47,48), AJ Woesearchaeota appear to have limited metabolic capabilities. It is therefore possible that these cells are partnered in a syntrophic relationship with other microbes on which they rely for energy or nutrients, as has been previously suggested (47,48).

The Woesearchaeotal MAG was one of several novel lineages in the AJ water column. From the deepest samples (95 m and 106 m), recovered MAGs included members of the sulfur-oxidizing SUP05 clade of *Thioglobaceae*, two members of the CPR phylum Patescibacteria, and three members of the phylum Marinimicrobia (SAR406), among others (Table 1). The middle water column (60 m) was also populated by underdescribed taxa, including several lineages of marine Thermoplasmatota and members of the bacterial phyla Planctomycetota and Myxococcota. Only eight out of nineteen MAGs from 60 m had identifiable close representatives in the GTDB, suggesting a high level of taxonomic novelty. Surprisingly, automatic binning did not recover a Thaumarchaeal MAG from the 60 m assembly despite the large fraction of amplicon SVs belonging to the ammonia-oxidizing genus *Nitrosopumilus*. However, a manually binned MAG could be phylogenomically placed in this lineage (Fig. S4) and contained the dominant SV from the amplicon data set, which comprised up to 86% of all *Nitrosopumilus* spp. amplicons. As with the Woesearchaeota, this low level of SV diversity implies population homogeneity. It remains to be determined if this homogeneity is driven by a potential dearth of ecological niches. Indeed, the AJ *Nitrosopumilus* MAG, at 725 Kbp, was estimated to be only 53-63% complete (Table 1, Table S3); its full biochemical potential therefore remains to be characterized. However, in contrast to recently described and putatively heterotrophic Thaumarchaeota (50,51), blue hole *Nitrosopumilus*, like all known members of this genus, contain *amo*CAB encoding ammonia monooxygenase and therefore likely contribute to nitrification in the blue hole.

The 60 m samples also contained six MAGs belonging to the archaeal phylum Thermoplasmatota. One of these (BH15) could only be placed in the family Thalassoarchaeaceae. The others included four (BH4, BH11, BH12, and BH13) from the Marine Group II lineage, which is one of the four major planktonic archaeal groups (52), and one from the less well-characterized Marine Group III lineage (BH3). The Marine Group II have recently been proposed as an order-level lineage, *Candidatus* Poseidoniales, containing two families delineating the current MGIIa and MGIIb clades (53). Most MGIIa members have been identified from the photic zone, while MGIIb sequences are largely limited to depths below 200 m, although there are exceptions (54). Notably, all AJ MGII MAGs belonged to the MGIIb lineage (Table 1). Several studies have identified genes related to low-oxygen metabolism including reduction of sulfate (55–57) and nitrate (53), suggesting these taxa are more likely to be adapted to microaerophilic or anaerobic conditions. We found a gene for thiosulfate reduction to sulfite (*phs*A) in two of the four MGIIb MAGs (Table 2), but no complete pathways for any forms of anaerobic respiration. Nevertheless, the Marine Group II lineage comprised over 25% (five out of nineteen) of the MAGs recovered from 60 m, suggesting this group as an important component to the hypoxic zone community.

Metabolic potential differentiated the middle and deep water column communities (Fig. 5, Fig. S6). Many of these functions could be linked to MAGs and represented a range of anaerobic metabolisms. Genes for dissimilatory sulfide and sulfite oxidation (or potentially reduction; *dsr*AB*, apr*AB), denitrification (*nar, nir, nor, nos*), and dissimilatory nitrate reduction to ammonia (DNRA, *nrf*) were enriched in deep anoxic samples, but also common in the hypoxic zone (60 m) (Table 2, Fig. 5). Other genes linked to MAGs included *aio*AB and *arr*A, encoding arsenite oxidase and arsenate reductase, respectively (Fig. S8). Reduction of arsenate to arsenite is thought to be an ancient metabolic pathway, originating before oxygenation of the Earth’s atmosphere and oceans (58). It has recently been proposed as an important microbial metabolic strategy in oxygen-deficient marine environments, potentially providing arsenite for use as an energy source by other microbes (19). Arsenic concentrations in Amberjack have not been measured. However, the detection of *aio*AB and *arr*A, of which the latter was also detected in unbinned contigs from the deep co-assembly (Fig. S8), as well as genes for arsenic detoxification (*ars*CM) in diverse MAGs (including in the *Nitrosopumilus* MAG described above), suggest a role for arsenic cycling in AJ. Remarkably, one MAG (BH24, *Rhodospirillales*) contained genes for several of these pathways, representing an unusual repertoire of respiration strategies (Table S4). Finally, one of the genes with the largest differences in frequency between 60 m and deep samples encodes an unknown protein previously observed in genomes of *Candidatus* Uhrbacteria, one of the groups within the CPR/Patescibacteria (Fig. S7). Two MAGs from this group, which is largely affiliated with freshwater and subsurface aquifer environments, were recovered from the deep samples, and may play as-yet undetermined ecological roles in this unusual microbiome. This hypothesis is supported by the low sequence identity of this protein to similar ones in NCBI.

## Conclusions

The Amberjack Hole water column is potentially highly stable, with complete turnover (driven either by wind or the sinking of cold water masses) unlikely. Thus, AJ microbiomes, particularly those in the deepest layers, potentially have limited connectivity to other marine communities. The extent to which such isolation explains the observed unique community composition or drives divergence of individual microbial lineages remains uncertain. Undoubtedly, the vertical transport, transformation, and stratification of redox active elements - notably sulfur, nitrogen, and iron - also play a significant role in structuring this unusual microbiome. The presumed stability of this community, its high metabolic diversity, and its dominance by understudied microbial taxa, highlight AJ as a model for detecting novel biogeochemical processes under low oxygen. Marine blue holes in general are potentially valuable sites for the study of microbial diversification and linked elemental cycling.

## Methods

### Sampling scheme

Sampling was conducted May 15-17 2019 and September 19 2019 using the vessels *R/V* William R. Mote and *R/V* Eugenie Clark. Amberjack Hole is located approximately 50 kilometers west of Sarasota, Florida, at 27.28748 N, −83.16139 W. One component of the research involved the deployment of an autonomous benthic lander to the bottom of the hole for 24-48 hours at a time for benthic electrochemistry measurements; these data are not included in this paper. Technical SCUBA divers who guided the lander into and out of the hole also collected water samples (see details below).

In both May and September, all water samples used for nutrient measurements and microbial DNA preservation were filtered immediately on board. Microbial DNA samples were stored on ice until the return to Mote Marine Laboratory, where they were stored at −20°C. Subsequent nutrient analyses described below were conducted at Mote except where noted.

### Water column physical parameters and dissolved oxygen

During the May sampling, an EXO2 sonde (YSI Inc/Xylem Inc, Yellow Springs, Ohio) was attached to the autonomous lander deployed for 24 hours to the bottom of the hole. The sensor provided extensive data from the surface (1-2 m) and the bottom position (106 m) but low-resolution readings from the water column due to rapid ascent and descent velocities. Dissolved oxygen, temperature, and salinity data were retrieved from the instrument and plotted in R.

During the fall sampling, an integrated SBE-19plus V2 CTD with an SBE43 DO sensor, WET Labs FLNTUrt chlorophyll and turbidity sensor, Satlantic cosine PAR sensor, and an SBE18 pH sensor (Sea-Bird Electronics Inc) was lowered twice by hand at a rate that provided high-resolution dissolved oxygen, turbidity, salinity, and density data on September 19 2019. CTD casts were performed the same day as water was sampled for microbial DNA and nutrients. Data were retrieved and plotted in R as above.

SBE data were processed using SBE Data Processing software after compensating for sensor thermal mass, sensor alignment, timing offset (the delay associated with pumped sensors, i.e. conductivity), and changes in instrument velocity (due to ship heave). Data were binned (~0.2 m increments) to smooth data variability. Processed and derived data included temperature (°C), potential temperature (°C), salinity (psu), water density (σ_T_ in kg/m^3^), chlorophyll *a* (μg/L based on relative fluorescence), turbidity (NTU), dissolved oxygen (mg/L), O_2_ saturation (%), pH, PAR, and Brunt-Väisälä frequency (N = stratification index). Manufacturer recommendations for calibration and service for all sensors were followed.

### Water column electrochemical measurements

Water samples (~10 m vertical resolution) were subject to solid-state Hg/Au voltammetric analyses (59) for measurement of the redox environment, i.e. O_2_, Mn(+II), organic-Fe(III) complexes, Fe(II), S(+II) (in the form of S_2_O_3_^−^), S(0), S_x_(-II), and ∑H_2_S (S^2−^, HS^−^, H_2_S). Briefly, samples collected from the water column at 10 m intervals were carefully transferred via a Tygon (formula 2375) transfer tube line into LDPE bottles while filling from bottom to top to minimize atmospheric oxygen contaminations. Sample bottles remained sealed until analyses and were stored at 4°C. Upon return to the lab (within 4-6 hours of collection), 20 mL of each sample was carefully pipetted into an electrochemical cell (Analytical Instrument Systems, Inc.) holding a custom fabricated Hg/Au amalgam microelectrode working electrode (60), a Pt counter electrode, and a fritted Ag/AgCl reference electrode (with 3 M KCl electrolyte solution). N_2_ was gently blown over the top of the solution to minimize mixing with the atmosphere. Within 5 minutes, samples were subject to a series of anodic square wave voltammograms (ASWVs), cathodic square wave voltammograms (CSWVs), and linear sweep voltammograms (LSVs) to quantify the above redox analytes (59,61). To analytically distinguish between S(0) and S(-II), which both react at the same potential at an Hg electrode (42), CSWV measurements were repeated after acidification with HCl (final pH < 3) and N_2_ sparging of the solution for two minutes to remove free ∑H_2_S. This same electrochemical speciation experiment was conducted a second time in separate aliquots that were filtered through 0.7 μm GFF filters directly in the N_2_-degassed electrochemical cell (this filter size was selected for future organic carbon analyses).

### Nutrient, iron, and DIC measurements

Water samples were collected for nutrients and included chlorophyll *a*, dissolved ammonium (NH_4_^+^), dissolved NO_x_ (nitrate (NO_3_) + nitrite (NO_2_)), dissolved orthophosphates (PO_4_^3−^), dissolved total and ferrous iron (Fe), dissolved inorganic carbon (DIC) and particulate carbon (C), nitrogen (N), and phosphorous (P). All samples were collected and immediately filtered once shipboard.

Chlorophyll *a* was measured according to EPA method 445.0 (62). Briefly, samples were filtered through a glass fiber filter until clogging, then stored in the dark until analyses. Prior to analyses, filters were sonicated in 90% acetone to extract chlorophyll from algal cells and then centrifuged for clarification. The fluorescence of the clarified extract was then measured using a fluorometer (Turner 10-AU Fluorometer) with special narrow bandpass filters at an excitation wavelength of 436 nm and an emission wavelength of 680 nm. Analytical quality assurance followed standard laboratory practices assessing precision and accuracy using sample replicates, container blanks, duplicate and spiked analyses with results meeting acceptable levels of precision and accuracy.

Samples for dissolved NH_4_^+^ and NO_x_ were filtered through Pall Supor (PES) 450 47 mm (0.45 μm pore size) membrane filters while samples for particulates were filtered through pre-combusted (450°C, 3h) 47 mm (0.7 μm pore size) Whatman GF/F filters. For particulate carbon and nitrogen, 200 mL was filtered per sample and filters were rinsed with acidified (10% HCl) filtered seawater to remove inorganic carbon. For particulate P, 500 mL was filtered. Samples were then stored on ice until their return to the lab and analyzed within 48 hours (dissolved NH_4_^+^ and NOx) or within 28 days (particulates). Analyses for dissolved nutrients followed colorimetric, segmented flow, autoanalyzer techniques on an AA3 with method reference and method detection limits (MDL) as follows: dissolved NH_4_^+^ ((63); 0.07 μM), dissolved NO_x_ ((64); 0.07 μM). Particulate phosphorus was analyzed according to (65,66) with in-house modifications for analysis on a segmented flow analyzer and MDLs of 0.03 μM. For particulate carbon and nitrogen, samples were analyzed on a Thermo FlashEA1 1112 Elemental Analyzer with MDLs of 0.2 and 0.1 μM, respectively.

To measure DIC, samples were immediately poisoned with HgCl_2_ and stored until analyses. DIC was analyzed (Apollo AS-C6 DIC Analyzer) following methods by (67). Accuracy and precision of the instrument was regularly monitored using Certified Reference Materials for Seawater CO_2_ Measurements (Dickson Laboratory, Scripps Institution of Oceanography, San Diego, CA; Batch #181 and 186).

Soluble orthophosphates (PO_4_^3−^) were measured spectrophotometrically using the molybdate-blue technique (68).

Finally, the speciation of iron was obtained by measuring Fe(II) by the ferrozine assay (69) in filtered samples before (dissolved Fe(II)) and after (total dissolved Fe) reduction by hydroxylamine (0.2 M) using a long waveguide spectrophotometric flow cell. Analytical quality assurance follows standard laboratory practices assessing precision and accuracy using sample replicates, container blanks, duplicate and spiked analyses with results meeting acceptable levels of precision and accuracy.

### Microbial DNA sampling and preservation

Water column samples were collected on May 15 and 17 2019 and September 19 2019. In May, 800 mL water was collected by divers from depths of 46 m, 61 m, and 106 m inside the hole. Sterilized Nalgene bottles filled with deionized water were taken down by the divers, opened at the appropriate depth, and DI water was replaced with ambient seawater. Niskin bottles deployed on the CTD rosette also collected 1.9 L samples from depths of 8 m, 15 m, and 23 m above the hole. In May and September, hand-cast Niskin bottles collected 1 L water from 0 m (surface), 30 m, 60 m, and 85 m (May), and from 0 m (surface) to 90 m at 10 m intervals, as well as 85 m and 95 m (September). Samples were filtered onto 0.22 μm Sterivex (MilliporeSigma) filters with a peristaltic pump and preserved with approximately 3 mL DNA/RNA stabilization buffer (25 mM sodium citrate, 10 mM EDTA, 5.3 M ammonium sulfate (pH 5.2)) and stored at −80°C or on dry ice until further processing at Georgia Tech. In September, unfiltered water from the following depths was also preserved (1.4 mL water, 150 μL PBS-buffered formaldehyde) for nucleic acid staining and microscopy and kept frozen at −20°C until processing: 30 m, 50 m, 60 m, 70 m, 80 m, 85 m, 90 m, 95 m.

### Sequencing library preparation

DNA was extracted from each Sterivex cartridge using a custom protocol as described in (70), except for the September surface (0 m) sample which was lost during sample transit to Georgia Tech. Briefly, cells were lysed by flushing out RNA stabilizing buffer and replacing it with lysis buffer (50mM Tris-HCl, 40 mM EDTA, 0.73 M sucrose) and lysozyme (2 mg in 40 mL of lysis buffer per cartridge), then incubating cartridges for 45 min at 37°C. Proteinase K was added and cartridges were resealed and incubated for 2 h at 55°C. The lysate was removed, and the DNA was extracted once with phenol:chloroform:isoamyl alcohol (25:24:1) and once with chloroform:isoamyl alcohol (24:1). Finally, DNA was concentrated by spin dialysis using Ultra-4 (100 kDA, Amicon) centrifugal filters. Yield was assessed using a Qubit 2.0 dsDNA high-sensitivity assay (Invitrogen, Carlsbad, CA).

Illumina MiSeq libraries were prepared by amplifying the V4 region of the 16S rRNA gene using the environmental DNA protocol adapted from (71). Briefly, amplicons were generated using Platinum^®^ PCR SuperMix (Life Technologies, Carlsbad, CA) with Earth Microbiome Project primers 515FB and 806RB appended with Illumina-specific adapters. Template DNA was diluted to approximately 5 ng/μL for all samples and PCRs were performed in 25-μL reactions using Platinum^®^ PCR SuperMix (Life Technologies) (22 μL), BSA (Invitrogen) (1 μL), and 0.5 μL each of forward and reverse primer (10 ng/L stock concentration) with 1 μL template DNA. The thermal cycling protocol consisted of 26 cycles with the following steps: denaturation at 98°C (30 s), followed by 30 cycles of denaturation at 98°C (5 s), primer annealing at 55°C (5 s) and primer extension at 72°C (8 s), followed by extension at 72°C for 1 minute. Negative control reactions were run using 1 μL milliQ water in place of DNA template. Amplicons were analyzed by gel electrophoresis to verify size (~400 bp, including barcodes and adaptor sequences) and purified using Diffinity RapidTip2 PCR purification tips (Diffinity Genomics, West Chester, PA). Amplicons from different samples were pooled at equimolar concentrations and sequenced using a paired-end Illumina MiSeq 500 cycle kit (2 × 250 bp) with 5% PhiX to increase read diversity.

Metagenomes were generated from four water column samples: two from May (60 m and 106 m) and two from September (60 m and 95 m). Libraries were prepared using the Illumina Nextera XT DNA library preparation kit (Illumina Inc., San Diego, CA) according to manufacturer’s instruction and run on a Bioanalyzer 2100 instrument (Agilent) using a high sensitivity DNA chip to determine library insert sizes. An equimolar mixture of the libraries (final loading concentration of 12 pM) was sequenced on an Illumina MiSeq instrument (School of Biological Sciences, Georgia Institute of Technology), using a MiSeq reagent v2 kit for 600 cycles (2 × 300 bp paired end protocol).

### Amplicon sequence data processing and analysis

Demultiplexed amplicon sequences are available in the Patin FigShare account: https://figshare.com/projects/Amberjack_Blue_Hole/85013. Raw sequences were run through the DADA2 algorithm (72) in QIIME2 (73) to assess sequences at sequence variant resolution, using the following command parameters: --p-trim-left-f 70 --p-trim-left-r 70 --p-trunc-len-f 150 --p-trunc-len-r 150. Resulting SVs were assigned taxonomy using the naïve Bayes classifier trained on the Silva 132 database (99% OTUs, 515F/805R sequence region).

The SV table with raw read counts of all water samples were exported from QIIME2 and transformed in R using the variance stabilizing transformation (vst) in the DESeq2 package (74). Tables were not rarefied to preserve maximum available information. Metadata including depth and SV taxonomy were imported and combined into a Phyloseq object (75,76). Alpha and beta diversity analyses were run in DivNet (77) at the SV level. Shannon and Simpson diversity results were extracted and plotted by depth grouping using ggplot2 (78). Beta diversity was assessed using the resulting Bray-Curtis distance matrix in a principal component analysis using the prcomp() function in R and visualized in a PCA using ggplot2. The distance matrix was also used to test for significant difference among samples by depth grouping and month using the adonis() function in the R vegan package (79) with the following command: adonis(formula = bc ~ Depth + Month, data = metadata, permutations = 999).

### Metagenomic sequence data processing and analysis

Metagenome sequences are available in the Patin FigShare account: https://figshare.com/projects/Amberjack_Blue_Hole/85013. Raw sequences from the four metagenomes (May 60 m, May 106 m, September 60 m, September 95 m) were trimmed and checked for quality and adapter contamination as described for the amplicons above. Quality controlled reads were run through MicrobeCensus (80) to generate genome equivalent (GE) values for each metagenome. To generate more even coverage distribution across the metagenomes, quality controlled reads were normalized using BBNorm, part of the open source BBMap package (https://sourceforge.net/projects/bbmap/).

Six assemblies were generated from the four water column metagenomes. Individual assemblies of each sample were generated using metaSPAdes (v3.14.0) (81). Co-assemblies for each depth pair (May 60 m with September 60 m, and May 106 m with September 95 m) were also generated using MEGAHIT (v1.2.9) (82) using a minimum contig length of 2500 bp. All assemblies were assessed for quality using metaQUAST (83) and annotated using Prokka (84) and the KEGG database.

For the latter, open reading frames were generated using Prodigal (85) and clustered using MeShClust (86) at 90% nucleotide identity. The longest sequence from each cluster was extracted using a custom Python script and these representative sequences were run against the KEGG ortholog profile HMM models (KOfams) using KofamScan with the ‘prokaryote’ database (87). The parameter ‘002D;f mapper’ was applied to provide only the most confident annotations (those assigned an individual KO). Orthologies were matched to their corresponding functions using a parsed version of the ‘kO00001.keg’ database text file (https://github.com/edgraham/GhostKoalaParser), which provides a three-tiered hierarchical categorization of each gene, referred to here as ‘Group’, ‘Subgroup1’, and ‘Subgroup2’. Sequence coverage of each gene was generated by mapping metagenomic short reads against each one using Magic-BLAST (88). The Magic-BLAST output was filtered to include only the best hit for each read and the read counts were normalized by GE value of the corresponding metagenome.

The normalized counts were used to run hierarchical clustering analyses in Python using the Seaborn ‘clustermap’ function. KofamScan outputs were grouped by the three hierarchical categories to generate heat maps at different levels of categorization. At the highest categorical level (‘Group’) the categories ‘Human Diseases,’ ‘Organismal Systems,’ ‘Cellular community – eukaryotes,’ and ‘Brite Hierarchies’ were removed before performing the cluster analysis. The first three groups are not relevant to microbial gene functions and the fourth provides a different hierarchical categorization scheme for the same annotations and was thus redundant. All cluster analyses were run using ward linkage and Euclidean distance methods.

Open reading frames generated from the SPAdes individual assemblies were run through the Hidden Markov Model search tool described in (89) and available at https://github.com/ShadeLab/PAPER_Dunivin_meta_arsenic/tree/master/HMM_search to query for genes involved in arsenic metabolism. HMM outputs were run through the data_preparation.R script (also available in the GitHub repository) to generate plots showing the quality distributions of hits to the *aio*A and *arr*A genes. The amino acid sequence of a potential sequence for the *arr*A gene extracted from BH30 (alignment fraction 89%, bitscore 237) was also queried against the *arr*A BLAST database from the same toolkit.

### Metagenome-assembled genomes

Genomic bins were generated from all individual assemblies as well as co-assemblies. For each co-assembly, short reads from both metagenomes were submitted to MaxBin to leverage contig co-variation patterns across samples. All bins were assessed for quality using CheckM (90) and anvi’o (v6.1) (91). Bins with a quality score greater 40 were retained for dereplication, with quality score calculated as completion – 5 × contamination (CheckM values). Dereplication was performed with dRep (92) with an ANI cutoff of 95% for secondary clustering and a 20% minimum pairwise overlap between genomes. For bins belonging to the same secondary cluster (ie, likely representing the same microbial population), the highest-quality was chosen as a representative. In cases where the qualities of two MAGs were within 1%, the completeness and redundancy values from anvi’o were applied to determine the higher quality bin. Further analyses were performed on these representative bins.

All dereplicated high-quality MAGs were assessed for taxonomy with anvi’o and the Genome Taxonomy Database toolkit (GTDB-Tk) (93,94). Each MAG was checked for the presence of rRNA genes using the anvi’o HMM models and one MAG from the 60M assembly was found to contain a eukaryotic 18S rRNA gene with an 80% match to a copepod sequence (*Calanus* sp.). This contig was removed from the co-assembly of origin and the process of binning, quality assessment, and de-replication was repeated.

All MAGs were queried for the presence of 16S rRNA gene amplicons by using the SVs as a query for a BLASTn analysis against the two co-assembly contigs. MAG contigs containing 100% identity matches across the entire SV length were identified, and the SV SILVA-based taxonomy was compared with the whole genome taxonomy assignment from GTDB.

All MAGs were functionally annotated using both Prokka and KofamScan as described above, without the read mapping step. The following genes were verified by running amino acid sequences against the NCBI-nr database using blastp: *pmo*A-*amo*A (K10944; KEGG), *pmo*B-*amo*B (K10955; KEGG), *pmo*C-*amo*C (K10956; KEGG), K23573 (DSPP, dentin sialophosphoprotein; KEGG), *pmo*A (Prokka), *pmo*B (Prokka), *aio*B (Prokka), *ars*C2 (Prokka), *ars*M (Prokka). To confirm the annotations of arsenic respiration and resistance genes, and to query other genes involved in arsenic metabolism, including the arsenate reductase gene *arr*A, all MAGs were run through the Hidden Markov Model search tool described in (89) and available at https://github.com/ShadeLab/PAPER_Dunivin_meta_arsenic/tree/master/HMM_search.

### Manual MAG refinement and generation

Six MAGs contained more than one 16S rRNA gene SV. In cases where the SVs within a MAG were distantly related to each other, any contigs containing a copy that was divergent from the genome-wide GTDB-TK taxonomic assessment were removed from the bin. Quality assessments were repeated on the edited MAG using CheckM and anvi’o, and if the quality score (as defined above) was over 5 points lower than the original MAG, the edited MAG was discarded. This happened in only one case (BH22); all other MAGs were kept in their edited form.

One of these edited MAGs was BH28, classified by GTDB as Patescibacteria but containing an SV classified as *Magnetospiraceae* (Alphaproteobacteria, Rhodospirillales) by SILVA. Conversely, BH24, which was classified by GTDB as Rhodospirillales, contained no 16S rRNA genes. The contig containing the *Magnetospiraceae* SV was removed from the Patescibacteria bin and added to BH24 based on its tetranucleotide clustering proximity to other contigs in that bin (as seen in the anvi’o interactive interface) and the matching taxonomy between the MAG and the SV.

The dominant *Nitrosopumilus* amplicon SV was not present in any of the MAGs generated from the 60 M samples. However, it was found on an unbinned contig in the 60M co-assembly. The anvi’o interactive interface was used to manually generate a bin containing this sequence, along with a 23S rRNA sequence with 99% identity to *Nitrosopumilus catalina*, and other contigs of similar tetranucleotide frequency, GC content, and coverage (BH19). This MAG was assessed for quality using CheckM and anvi’o.

All MAGs were run through GTDB using the ‘classify_wf’ (for taxonomic classification) and ‘ani_rep’ (for closest relative by ANI) commands in GTDB-Tk (93), and through Prokka (84) and KofamScan (87) for functional annotation. Closest relatives were determined using a minimum 65% alignment fraction; if no reference genome exceeded this minimum there was no result.

All MAG sequences are available in the Patin FigShare account: https://figshare.com/projects/Amberjack_Blue_Hole/85013.

### Phylogenomic analyses

Phylogenomic trees were generated for the MAGs BH19 (*Nitrosopumilus*), BH20 (SUP05 clade, *Thioglobaceae*), and BH21 (Woesearchaeota). In each case, genomes with the same or similar taxonomy as the MAG were downloaded from the NCBI Assembly database and run through anvi’o for quality assessment using anvi-estimate-genome-completeness. High-quality MAGs (quality score > 40 as defined above) were retained (Table S4) and single-copy genes were extracted using the anvi’o script anvi-get-sequences-for-hmm-hits. The two archaeal MAGs (BH19 and BH21) were run against the ‘Archaea_76’ HMM database and the bacterial MAG (BH20) was run against the ‘Bacteria_71’ HMM database. Genes occurring in most or all MAGs were concatenated and aligned (see Table S4 for taxon-specific minimum gene occurrences). The alignment was run through RAxML (95) with optimization of substitution rates and a GAMMA model of rate heterogeneity (“PROTGAMMA” substitution model) with 999 bootstraps for a maximum likelihood phylogeny. The resulting .tre files were uploaded to the interactive Tree of Life website (96) (itol.embl.de) and labeled with their strain or sample of origin.

### Cell staining and microscopy

Duplicate samples from the following depths were processed for microscopy: 30 m, 50 m, 70 m, 80 m, and 95 m. Samples preserved in PBS-buffered formaldehyde (1.55 mL total) were combined with 2.0 mL milliQ water and filtered onto 0.2 um GTBP filters (Millipore) using vacuum filtration. Filters were dried for approximately 20 minutes and incubated in the dark on ice with 50 μL DAPI (0.2 μg/mL). Filters were then rinsed in milliQ water and 100% ethanol, dried for approximately 10 minutes, and placed on a microscope slide with one drop of Citifluor (Electron Microscopy Sciences). Slides were visualized with a Zeiss Axio Observer D1 confocal epifluorescence microscope using a DAPI filter (Zeiss filter set 49). Between twenty and twenty-five photographs were taken in a grid pattern from each slide and counted in ImageJ (97) using a custom script. The average value of cell counts from all photos for each sample was plotted in Microsoft Excel.

## Table Legends

Table 1. All de-replicated, high-quality metagenome-assembled genomes (MAGs) from the 60 m and deep co-assemblies, as well as one MAG from the September 95 m individual assembly. Taxonomy assessed at the whole-genome level and nearest relative (GTDB) are provided. If applicable, taxonomy according to the associated 16S rRNA gene (SILVA) is provided with additional higher-order classification in parentheses. Amplicon frequencies for corresponding SVs are provided for each sample. One MAG contained two SVs (*, see Table S3) and here the frequencies for the first SV are provided.

Table 2. Taxonomy and functional annotations identified in each MAG. The biochemical process associated with each gene is provided by headers in bold.

Table S1. Amberjack Hole water column sample metadata and amplicon sequencing results. Processed reads are those that were quality-filtered, trimmed, and run through DADA2.

Table S2. Metagenome assembly data including sample source, pre- and post-quality filtered reads, and assembly statistics for both single and co-assemblies.

Table S3. Metagenome-assembled genomes (MAGs) and their associated samples, quality metrics, and assigned taxonomy.

Table S4. Metagenome-assembled genomes (MAGs) assessed in the phylogenomic trees (Fig. 4, Fig. S4, Fig. S5) including the Amberjack MAG of interest and the source of all other MAGs included in the phylogeny.

## Acknowledgments

We thank the captains of the *R/V* William R. Mote and *R/V* Eugenie Clark for their support and professionalism, and staff at Mote Marine Lab for their work in sample collection and processing. We are grateful to Dr. Navid Constantinou for his help modeling the wind speeds required for turnover of the water column.

## References

1. Mylroie JE, Carew JL, Moore AI. Blue Holes: Definition and Genesis. Carbonates and Evaporates. 1995;10(2):225–33.

2. Canganella F, Bianconi G, Kato C, Gonzalez J. Microbial ecology of submerged marine caves and holes characterized by high levels of hydrogen sulphide. In: Reviews in Environmental Science and Biotechnology. 2007.

3. Gischler E, Shinn EA, Oschmann W, Fiebig J, Buster NA. A 1500-Year Holocene Caribbean Climate Archive from the Blue Hole, Lighthouse Reef, Belize. J Coast Res. 2008;246:1495–505.

4. Pohlman JW. The biogeochemistry of anchialine caves: Progress and possibilities. Hydrobiologia. 2011;677:33–51.

5. Seymour JR, Humphreys WF, Mitchell JG. Stratification of the microbial community inhabiting an anchialine sinkhole. Aquat Microb Ecol. 2007;

6. Garman KM, Rubelmann H, Karlen DJ, Wu T, Garey JR. Comparison of an inactive submarine spring with an active nearshore anchialine spring in Florida. Hydrobiologia. 2011;

7. Gonzalez BC, Iliffe TM, Macalady JL, Schaperdoth I, Kakuk B. Microbial hotspots in anchialine blue holes: Initial discoveries from the Bahamas. Hydrobiologia. 2011;677(1):149–56.

8. Davis MC, Garey JR. Microbial function and hydrochemistry within a stratified anchialine sinkhole: A window into coastal aquifer interactions. Water. 2018;10(8):972.

9. Yao P, Wang XC, Bianchi TS, Yang ZS, Fu L, Zhang XH, et al. Carbon Cycling in the World’s Deepest Blue Hole. J Geophys Res Biogeosciences. 2020;

10. He H, Fu L, Liu Q, Fu L, Bi N, Yang Z, et al. Community Structure, Abundance and Potential Functions of Bacteria and Archaea in the Sansha Yongle Blue Hole, Xisha, South China Sea. Front Microbiol. 2019;10:2404.

11. He P, Xie L, Zhang X, Li J, Lin X, Pu X, et al. Microbial Diversity and Metabolic Potential in the Stratified Sansha Yongle Blue Hole in the South China Sea. Sci Rep. 2020;10:5949.

12. DeWitt D. Submarine Springs and other Karst Features in Offshore Waters of the Gulf of Mexico and Tampa Bay, Southwest Florida Water Management District. 2003.

13. Hughes JD, Vacher HL, Sanford WE. Three-dimensional flow in the Florida platform: Theoretical analysis of Kohout convection at its type locality. Geology. 2007;35(7):663–6.

14. Hu C, Muller-Karger FE, Swarzenski PW. Hurricanes, submarine groundwater discharge, and Florida’s red tides. Geophys Res Lett. 2006;33(11):L11601.

15. Smith CG, Swarzenski PW. An investigation of submarine groundwater-borne nutrient fluxes to the west Florida shelf and recurrent harmful algal blooms. Limnol Oceanogr. 2012;57(2):471–85.

16. Walsh JJ, Jolliff JK, Darrow BP, Lenes JM, Milroy SP, Remsen A, et al. Red tides in the Gulf of Mexico: Where, when, and why? J Geophys Res. 2006;111(C11003):1–46.

17. Weisberg RH, Liu Y, Lembke C, Lu C, Hubbard C, Garrett M. The coastal ocean circulation influence on the 2018 West Florida Shelf K. brevis red tide bloom. J Geophys Res Ocean. 2019;124(4):2501–12.

18. Vargo GA, Heil CA, Fanning KA, Dixon LK, Neely MB, Lester K, et al. Nutrient availability in support of Karenia brevis blooms on the central West Florida Shelf: What keeps Karenia blooming? Cont Shelf Res. 2008;28(1):73–98.

19. Saunders JK, Fuchsman CA, McKay C, Rocap G. Complete arsenic-based respiratory cycle in the marine microbial communities of pelagic oxygen-deficient zones. Proc Natl Acad Sci U S A. 2019;116(20):9925–30.

20. Callbeck CM, Lavik G, Ferdelman TG, Fuchs B, Gruber-Vodicka HR, Hach PF, et al. Oxygen minimum zone cryptic sulfur cycling sustained by offshore transport of key sulfur oxidizing bacteria. Nat Commun. 2018;9:1729.

21. Garcia-Robledo E, Padilla CC, Aldunate M, Stewart FJ, Ulloa O, Paulmier A, et al. Cryptic oxygen cycling in anoxic marine zones. Proc Natl Acad Sci U S A. 2017;114(31):8319–1824.

22. Sun X, Kop LFM, Lau MCY, Frank J, Jayakumar A, Lücker S, et al. Uncultured Nitrospina-like species are major nitrite oxidizing bacteria in oxygen minimum zones. ISME J. 2019;13:2391–402.

23. Tsementzi D, Wu J, Deutsch S, Nath S, Rodriguez-R LM, Burns AS, et al. SAR11 bacteria linked to ocean anoxia and nitrogen loss. Nature. 2016;536:179–83.

24. Thamdrup B, Steinsdóttir HGR, Bertagnolli AD, Padilla CC, Patin N V., Garcia-Robledo E, et al. Anaerobic methane oxidation is an important sink for methane in the ocean’s largest oxygen minimum zone. Limnol Oceanogr. 2019;64(6):2569–85.

25. Breitburg D, Levin LA, Oschlies A, Grégoire M, Chavez FP, Conley DJ, et al. Declining oxygen in the global ocean and coastal waters. Science (80-). 2018;359(6371):eaam7240.

26. Baker BJ, De Anda V, Seitz KW, Dombrowski N, Santoro AE, Lloyd KG. Diversity, ecology and evolution of Archaea. Nat Microbiol. 2020;5:887–900.

27. Thiel V, Costas AMG, Fortney NW, Martinez JN, Tank M, Roden EE, et al. “Candidatus Thermonerobacter thiotrophicus,” a non-phototrophic member of the Bacteroidetes/Chlorobi with dissimilatory sulfur metabolism in hot spring mat communities. Front Microbiol. 2019;9:3159.

28. Helly JJ, Levin LA. Global distribution of naturally occurring marine hypoxia on continental margins. Deep Res Part I Oceanogr Res Pap. 2004;51(9):1159–68.

29. Wyrtki K. The oxygen minima in relation to ocean circulation. Deep Res Oceanogr Abstr. 1962;9(1–2):11–23.

30. Thamdrup B, Dalsgaard T, Revsbech NP. Widespread functional anoxia in the oxygen minimum zone of the Eastern South Pacific. Deep Res Part I Oceanogr Res Pap. 2012;65:36–45.

31. Garman KM, Garey JR. The transition of a freshwater karst aquifer to an anoxic marine system. Estuaries. 2005;28:686–93.

32. Hughes JD, Vacher HL, Sanford WE. Temporal response of hydraulic head, temperature, and chloride concentrations to sea-level changes, Floridan aquifer system, USA. Hydrogeol J. 2009;17(4):793–815.

33. Simpson JH. The shelf-sea fronts: implications of their existence and behaviour. Philos Trans R Soc London Ser A, Math Phys Sci. 1981;302(1472):531–46.

34. Luther GW, Findlay AJ, MacDonald DJ, Owings SM, Hanson TE, Beinart RA, et al. Thermodynamics and kinetics of sulfide oxidation by oxygen: a look at inorganically controlled reactions and biologically mediated processes in the environment. Front Microbiol. 2011;2(62):1–9.

35. Ghosh W, Dam B. Biochemistry and molecular biology of lithotrophic sulfur oxidation by taxonomically and ecologically diverse bacteria and archaea. FEMS Microbiol Rev. 2009;33(6):999–1043.

36. Wright JJ, Konwar KM, Hallam SJ. Microbial ecology of expanding oxygen minimum zones. Nat Rev Microbiol. 2012;10:381–94.

37. Walsh DA, Zaikova E, Howes CG, Song YC, Wright JJ, Tringe SG, et al. Metagenome of a versatile chemolithoautotroph from expanding oceanic dead zones. Science (80-). 2009;326(5952):578–82.

38. Hawley AK, Brewer HM, Norbeck AD, Pasǎ-Tolić L, Hallam SJ. Metaproteomics reveals differential modes of metabolic coupling among ubiquitous oxygen minimum zone microbes. Proc Natl Acad Sci U S A. 2014;111(31):11395–400.

39. Bertagnolli AD, Stewart FJ. Microbial niches in marine oxygen minimum zones. Nat Rev Microbiol. 2018;16:723–9.

40. Wirsen CO, Sievert SM, Cavanaugh CM, Molyneaux SJ, Ahmad A, Taylor LT, et al. Characterization of an autotrophic sulfide-oxidizing marine Arcobacter sp. that produces filamentous sulfur. Appl Environ Microbiol. 2002;68(1):316–25.

41. Rozan TF, Theberge SM, Luther G. Quantifying elemental sulfur (S0), bisulfide (HS-) and polysulfides (S(x)2-) using a voltammetric method. Anal Chim Acta. 2000;415(1–2):175–84.

42. Luther GW, Glazer BT, Hohmann L, Popp JI, Tailefert M, Rozan TF, et al. Sulfur speciation monitored in situ with solid state gold amalgam voltammetric microelectrodes: Polysulfides as a special case in sediments, microbial mats and hydrothermal vent waters. J Environ Monit. 2001;3:61–6.

43. Heylen K, Vanparys B, Wittebolle L, Verstraete W, Boon N, De Vos P. Cultivation of denitrifying bacteria: Optimization of isolation conditions and diversity study. Appl Environ Microbiol. 2006;72(4):2637–43.

44. Taillefert M, Bono AB, Luther GW. Reactivity of Freshly Formed Fe(III) in Synthetic Solutions and (Pore)Waters: Voltammetric Evidence of an Aging Process. Environ Sci Technol. 2000;34(11):2169–77.

45. Barco RA, Emerson D, Sylvan JB, Orcutt BN, Jacobson Meyers ME, Ramírez GA, et al. New insight into microbial iron oxidation as revealed by the proteomic profile of an obligate iron-oxidizing chemolithoautotroph. Appl Environ Microbiol. 2015;81(17):5927–37.

46. Garber AI, Nealson KH, Okamoto A, McAllister SM, Chan CS, Barco RA, et al. FeGenie: A Comprehensive Tool for the Identification of Iron Genes and Iron Gene Neighborhoods in Genome and Metagenome Assemblies. Front Microbiol. 2020;11:37.

47. Castelle CJ, Wrighton KC, Thomas BC, Hug LA, Brown CT, Wilkins MJ, et al. Genomic expansion of domain archaea highlights roles for organisms from new phyla in anaerobic carbon cycling. Curr Biol. 2015;25(6):690–701.

48. Liu X, Li M, Castelle CJ, Probst AJ, Zhou Z, Pan J, et al. Insights into the ecology, evolution, and metabolism of the widespread Woesearchaeotal lineages. Microbiome. 2018;6:102.

49. Ortiz-Alvarez R, Casamayor EO. High occurrence of Pacearchaeota and Woesearchaeota (Archaea superphylum DPANN) in the surface waters of oligotrophic high-altitude lakes. Environ Microbiol Rep. 2016;8(2):210–7.

50. Aylward FO, Santoro AE. Heterotrophic Thaumarchaea with Small Genomes Are Widespread in the Dark Ocean. mSystems. 2020;5(3):e00415–20.

51. Reji L, Francis CA. Metagenome-assembled genomes reveal unique metabolic adaptations of a basal marine Thaumarchaeota lineage. ISME J. 2020;14:2105–2115.

52. Santoro AE, Richter RA, Dupont CL. Planktonic Marine Archaea. Ann Rev Mar Sci. 2019;11:131–58.

53. Rinke C, Rubino F, Messer LF, Youssef N, Parks DH, Chuvochina M, et al. A phylogenomic and ecological analysis of the globally abundant Marine Group II archaea (Ca. Poseidoniales ord. nov.). ISME J. 2019;13:663–75.

54. Pereira O, Hochart C, Auguet JC, Debroas D, Galand PE. Genomic ecology of Marine Group II, the most common marine planktonic Archaea across the surface ocean. Microbiol Open. 2019;8(9):e00852.

55. Martin-Cuadrado AB, Garcia-Heredia I, Moltó AG, López-Úbeda R, Kimes N, López-García P, et al. A new class of marine Euryarchaeota group II from the mediterranean deep chlorophyll maximum. ISME J. 2015;9:1619–34.

56. Moreira D, Rodríguez-Valera F, López-García P. Analysis of a genome fragment of a deep-sea uncultivated Group II euryarchaeote containing 16S rDNA, a spectinomycin-like operon and several energy metabolism genes. Environ Microbiol. 2004;6(9):959–69.

57. Martin-Cuadrado AB, Rodriguez-Valera F, Moreira D, Alba JC, Ivars-Martínez E, Henn MR, et al. Hindsight in the relative abundance, metabolic potential and genome dynamics of uncultivated marine archaea from comparative metagenomic analyses of bathypelagic plankton of different oceanic regions. ISME J. 2008;2:865–86.

58. Sforna MC, Philippot P, Somogyi A, Van Zuilen MA, Medjoubi K, Schoepp-Cothenet B, et al. Evidence for arsenic metabolism and cycling by microorganisms 2.7 billion years ago. Nat Geosci. 2014;7:811–5.

59. Luther GW, Glazer BT, Ma S, Trouwborst RE, Moore TS, Metzger E, et al. Use of voltammetric solid-state (micro)electrodes for studying biogeochemical processes: Laboratory measurements to real time measurements with an in situ electrochemical analyzer (ISEA). Mar Chem. 2008;108(3–4):221–35.

60. Brendel PJ, Luther GW. Development of a Gold Amalgam Voltammetric Microelectrode for the Determination of Dissolved Fe, Mn, O2, and S(-II) in Porewaters of Marine and Freshwater Sediments. Environ Sci Technol. 1995;29(3):751–61.

61. Tercier-Waeber M Lou, Taillefert M. Remote in situ voltammetric techniques to characterize the biogeochemical cycling of trace metals in aquatic systems. J Environ Monit. 2008;10(1):30–54.

62. Arar EJ, Collins GB. Method 445.0 In Vitro Determination of Chlorophyll a and Pheophytin a in Marine and Freshwater Algae by Fluorescence. 1997.

63. Bran+Luebbe/Seal. Ammonia in water and seawater. Method No G-171-96 Rev 10. 2005;Norderstedt, Germany.

64. Bran+Luebbe/Seal. Nitrate and nitrite in water and seawater; total nitrogen in persulfate digests. Method No G-172-96 Rev 10. 2010;Norderstedt, Germany.

65. Solórzano L, Sharp JH. Determination of total dissolved phosphorus and particulate phosphorus in natural waters. Limnol Oceanogr. 1980;25(4):754–8.

66. Solórzano L, Sharp JH. Determination of total dissolved nitrogen in natural waters. Limnol Oceanogr. 1980;25(4):751–4.

67. Dickson AG, Sabine CL, Christian JR. Guide to best practices for ocean CO2 measurements. PICES Spec Publ 3. 2007;

68. Murphy J, Riley JP. A modified single solution method for the determination of phosphate in natural waters. Anal Chim Acta. 1962;27:31–6.

69. Stookey LL. Ferrozine-A New Spectrophotometric Reagent for Iron. Anal Chem. 1970;42(7):779–81.

70. Padilla CC, Bertagnolli AD, Bristow LA, Sarode N, Glass JB, Thamdrup B, et al. Metagenomic binning recovers a transcriptionally active gammaproteobacterium linking methanotrophy to partial denitrification in an anoxic oxygen minimum zone. Front Mar Sci. 2017;4:23.

71. Kozich JJ, Westcott SL, Baxter NT, Highlander SK, Schloss PD. Development of a dual-index sequencing strategy and curation pipeline for analyzing amplicon sequence data on the MiSeq Illumina sequencing platform. Appl Environ Microbiol. 2013;79(17):5112–20.

72. Callahan BJ, McMurdie PJ, Rosen MJ, Han AW, Johnson AJA, Holmes SP. DADA2: High-resolution sample inference from Illumina amplicon data. Nat Methods. 2016;13:581–3.

73. Bolyen E, Rideout JR, Dillon MR, Bokulich NA, Abnet CC, Al-Ghalith GA, et al. Reproducible, interactive, scalable and extensible microbiome data science using QIIME 2. Nat Biotechnol. 2019;37:852–7.

74. Love MI, Huber W, Anders S. Moderated estimation of fold change and dispersion for RNA-seq data with DESeq2. Genome Biol. 2014;15:550.

75. McMurdie PJ, Holmes S. Phyloseq: An R Package for Reproducible Interactive Analysis and Graphics of Microbiome Census Data. PLoS One. 2013;8(4):e61217.

76. McMurdie PJ, Holmes S. Waste Not, Want Not: Why Rarefying Microbiome Data Is Inadmissible. PLoS Comput Biol. 2014;10(4):e1003531.

77. Willis AD, Martin BD. DivNet: Estimating diversity in networked communities. bioRxiv. 2018;

78. Wickham H. ggplot2: Elegant Graphics for Data Analysis. New York: Springer-Verlag; 2016.

79. Oksanen J, Blanchet FG, Friendly M, Kindt R, Legendre P, Mcglinn D, et al. vegan: Community Ecology Package [Internet]. Community ecology package. 2019. p. R package version 2.5-5. Available from: https://cran.r-project.org/package=vegan

80. Nayfach S, Pollard KS. Average genome size estimation improves comparative metagenomics and sheds light on the functional ecology of the human microbiome. Genome Biol. 2015;16:51.

81. Nurk S, Meleshko D, Korobeynikov A, Pevzner PA. MetaSPAdes: A new versatile metagenomic assembler. Genome Res. 2017;27(5):824–34.

82. Li D, Liu CM, Luo R, Sadakane K, Lam TW. MEGAHIT: An ultra-fast single-node solution for large and complex metagenomics assembly via succinct de Bruijn graph. Bioinformatics. 2015;31(10):1674–6.

83. Mikheenko A, Saveliev V, Gurevich A. MetaQUAST: Evaluation of metagenome assemblies. Bioinformatics. 2016;32(7):1088–90.

84. Seemann T. Prokka: Rapid prokaryotic genome annotation. Bioinformatics. 2014;30(14):2068–9.

85. Hyatt D, Chen GL, LoCascio PF, Land ML, Larimer FW, Hauser LJ. Prodigal: Prokaryotic gene recognition and translation initiation site identification. BMC Bioinformatics. 2010;11:119.

86. James BT, Luczak BB, Girgis HZ. MeShClust: an intelligent tool for clustering DNA sequences. Nucleic Acids Res. 2018;46(14):e83.

87. Aramaki T, Blanc-Mathieu R, Endo H, Ohkubo K, Kanehisa M, Goto S, et al. KofamKOALA: KEGG Ortholog assignment based on profile HMM and adaptive score threshold. Bioinformatics. 2020;

88. Boratyn GM, Thierry-Mieg J, Thierry-Mieg D, Busby B, Madden TL. Magic-BLAST, an accurate RNA-seq aligner for long and short reads. BMC Bioinformatics. 2019;

89. Dunivin TK, Yeh SY, Shade A. A global survey of arsenic-related genes in soil microbiomes. BMC Biol. 2019;17:45.

90. Parks DH, Imelfort M, Skennerton CT, Hugenholtz P, Tyson GW. CheckM: Assessing the quality of microbial genomes recovered from isolates, single cells, and metagenomes. Genome Res. 2015;25(7):1043–55.

91. Eren AM, Esen OC, Quince C, Vineis JH, Morrison HG, Sogin ML, et al. Anvi’o: An advanced analysis and visualization platformfor’omics data. PeerJ. 2015;3:e1319.

92. Olm MR, Brown CT, Brooks B, Banfield JF. DRep: A tool for fast and accurate genomic comparisons that enables improved genome recovery from metagenomes through de-replication. ISME J. 2017;11(12):2864–8.

93. Chaumeil P-A, Mussig AJ, Hugenholtz P, Parks DH. GTDB-Tk: a toolkit to classify genomes with the Genome Taxonomy Database. Bioinformatics. 2019;36(6):1925–7.

94. Parks DH, Chuvochina M, Waite DW, Rinke C, Skarshewski A, Chaumeil PA, et al. A standardized bacterial taxonomy based on genome phylogeny substantially revises the tree of life. Nat Biotechnol. 2018;36:996–1004.

95. Stamatakis A. RAxML version 8: A tool for phylogenetic analysis and post-analysis of large phylogenies. Bioinformatics. 2014;30(9):1312–1313.

96. Letunic I, Bork P. Interactive Tree of Life (iTOL) v4: Recent updates and new developments. Nucleic Acids Res. 2019;47(W1):W256–W259.

97. Schindelin J, Arganda-Carreras I, Frise E, Kaynig V, Longair M, Pietzsch T, et al. Fiji: An open-source platform for biological-image analysis. Nat Methods. 2012;9:676–682.

