## Supplementary figures and images for "Gulf of Mexico blue hole harbors high levels of novel microbial lineages"

### Supplemental Figure 1

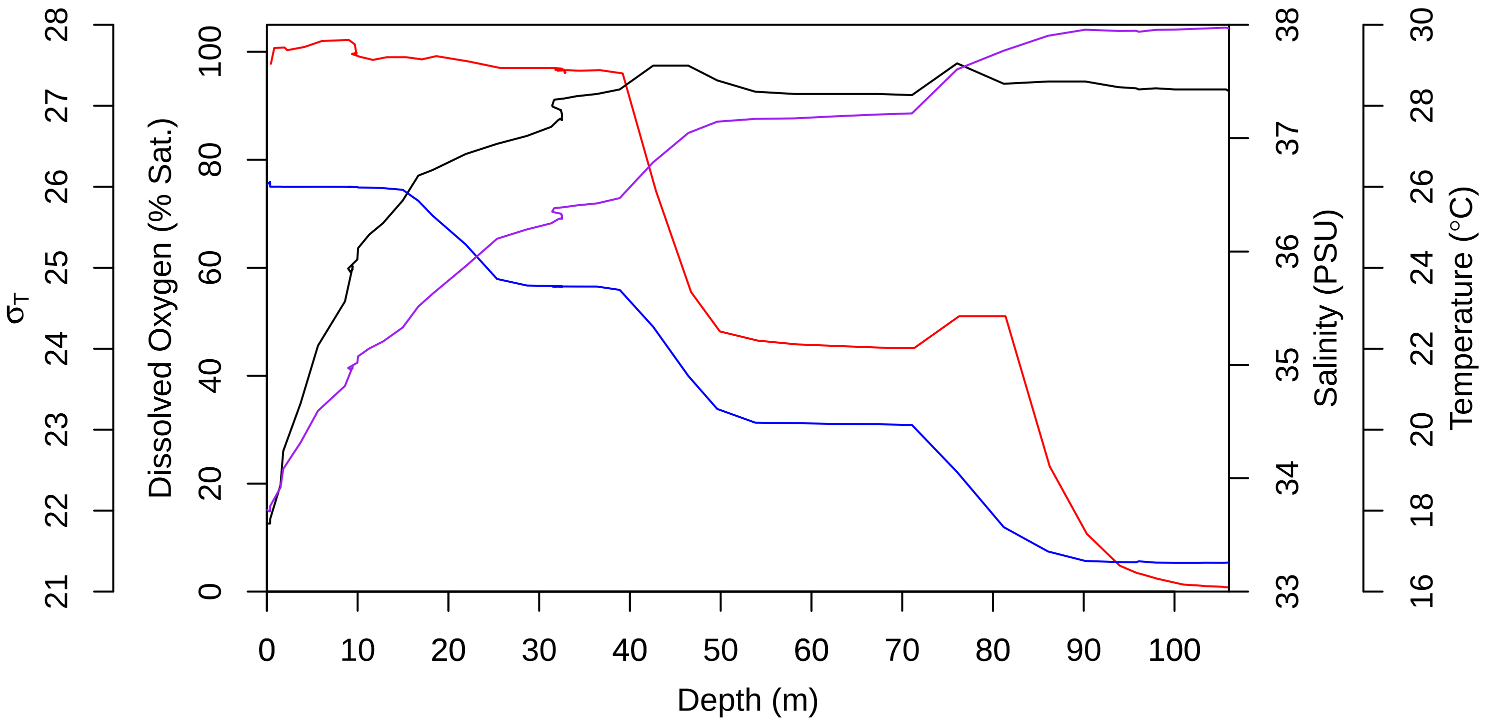

### Supplemental Figure 2

Figure S2. Chlorophyll a and particulate nutrient profiles from May (A) and September (B) 2019.

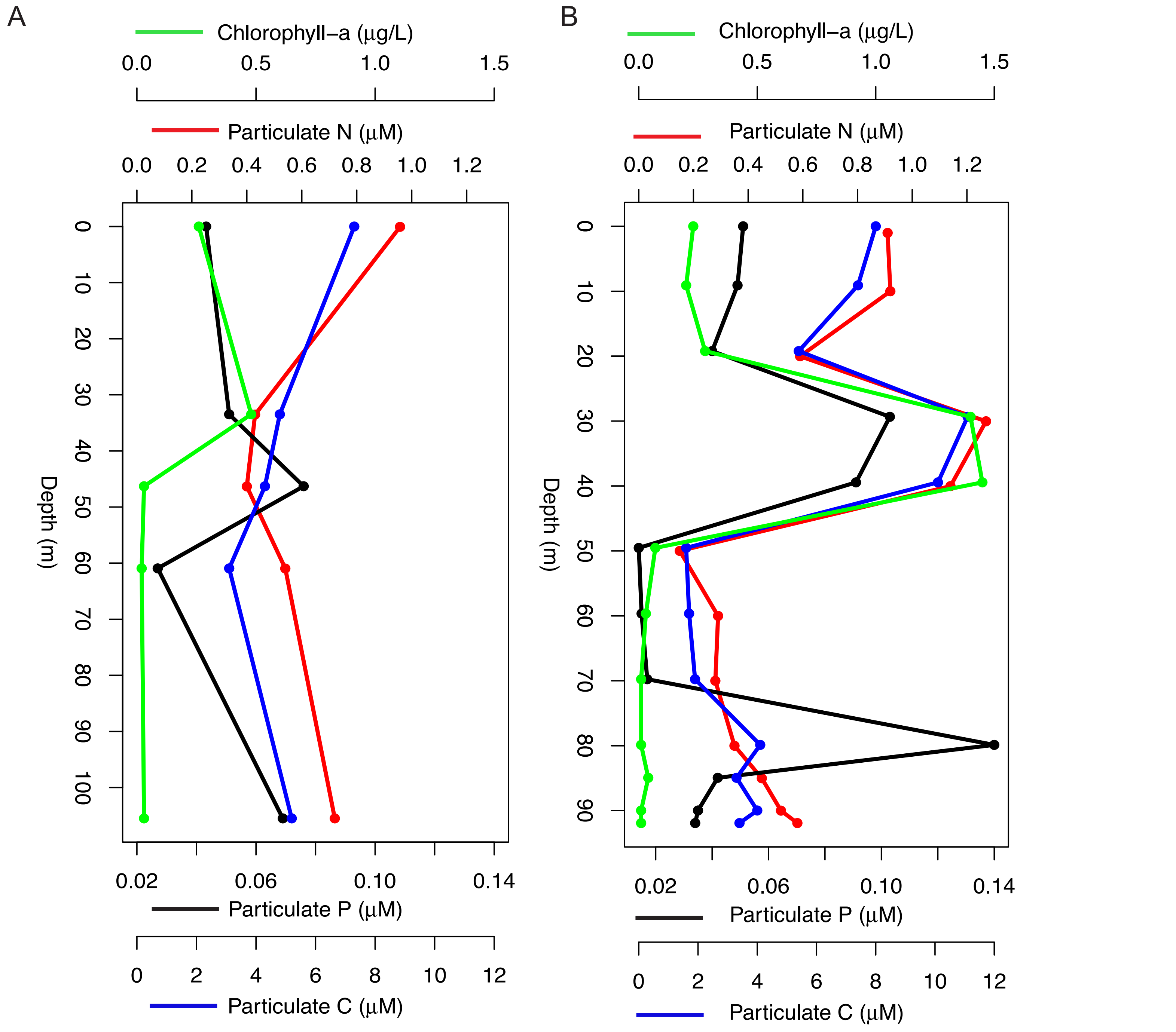

### Supplemental Figure 4

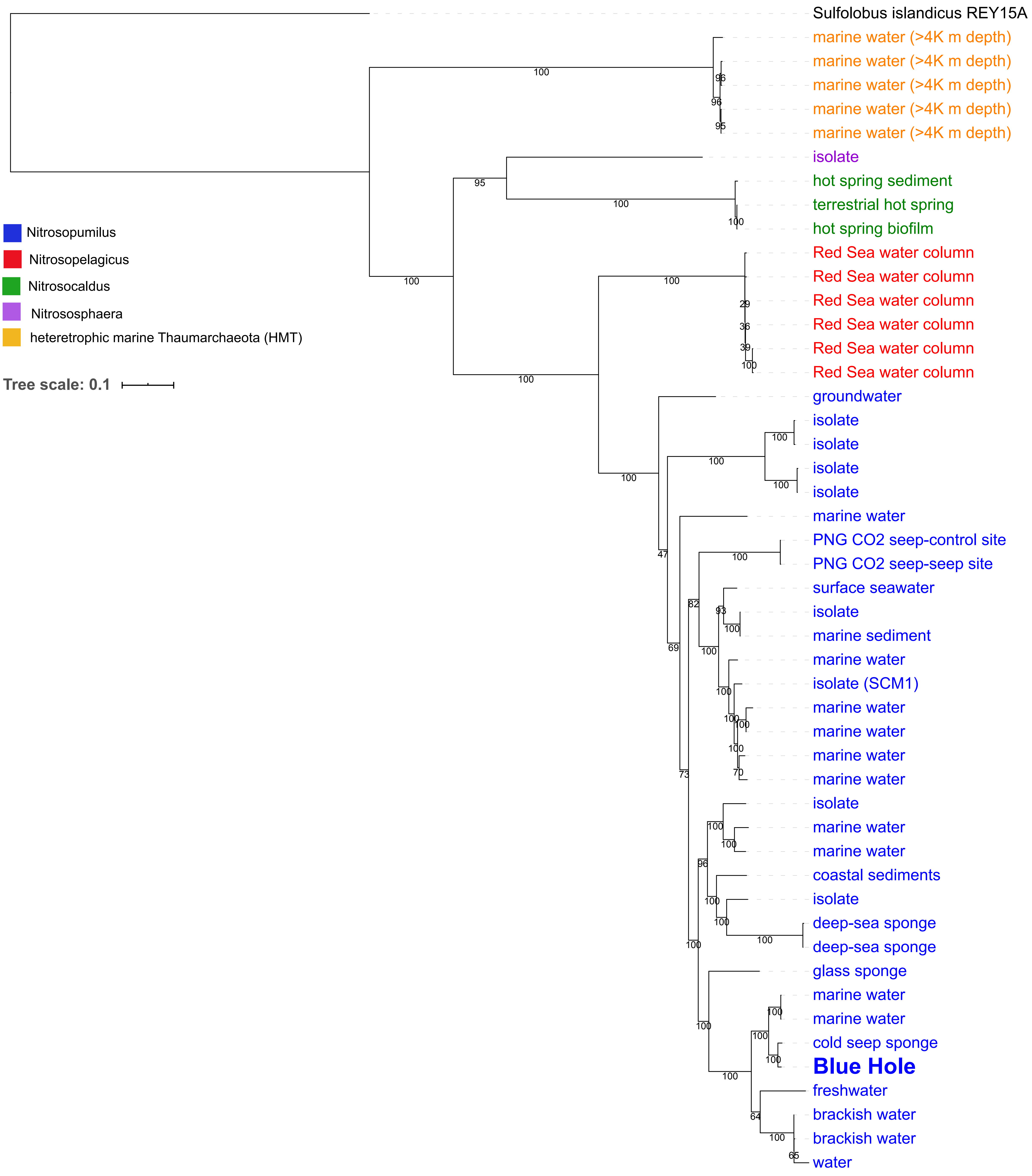

### Supplemental Figure 5

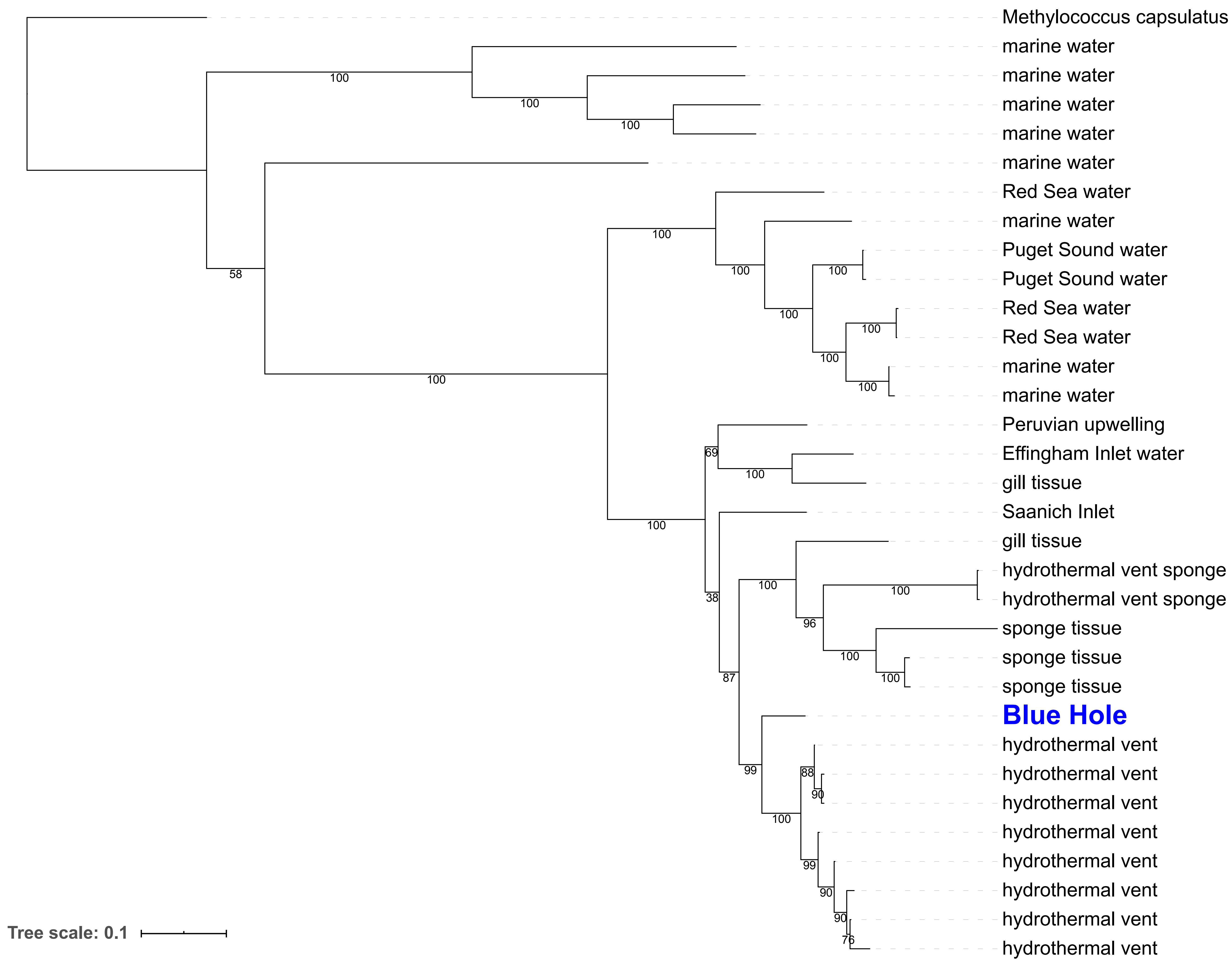

### Supplemental Figure 6

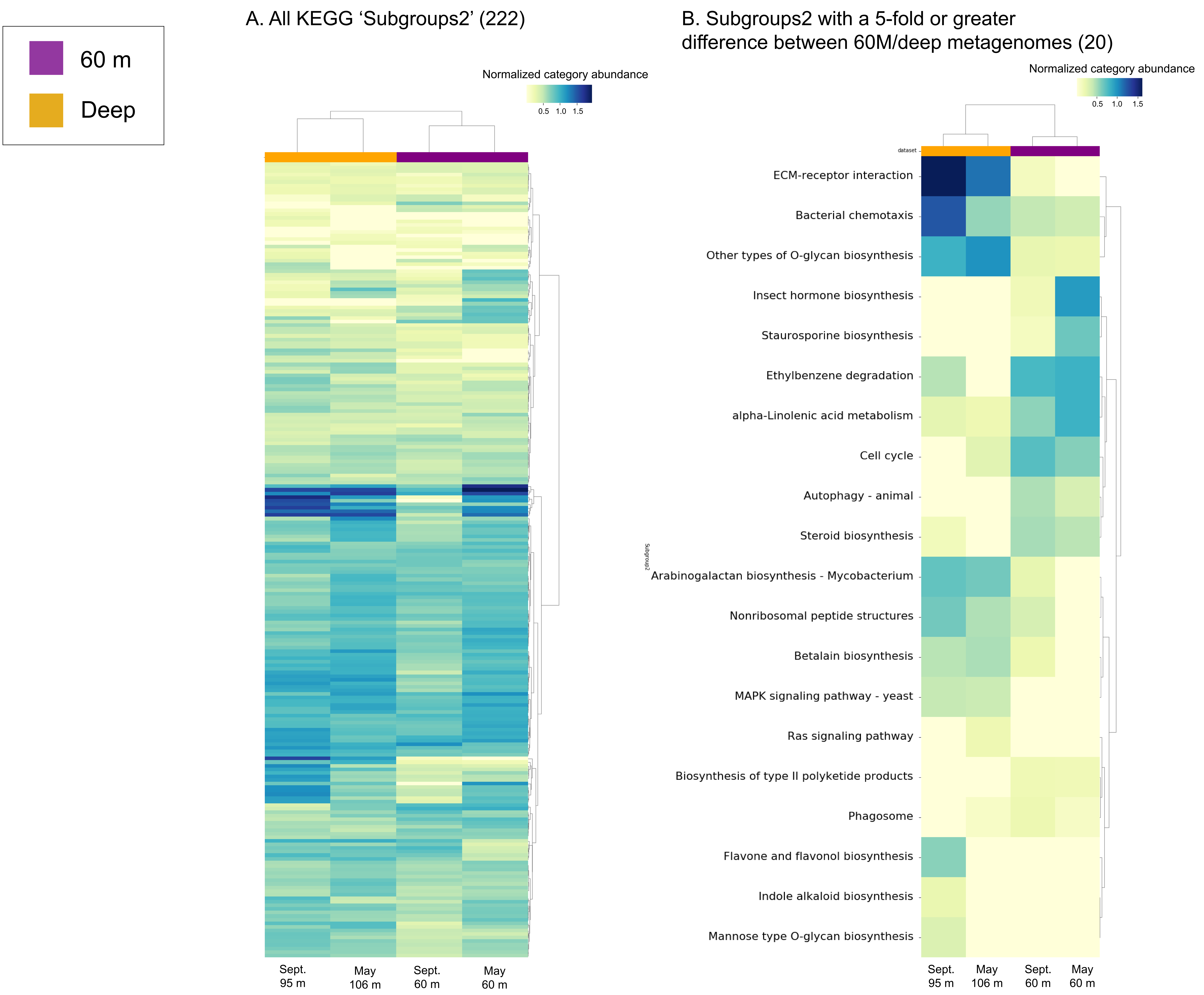

### Supplemental Figure 8

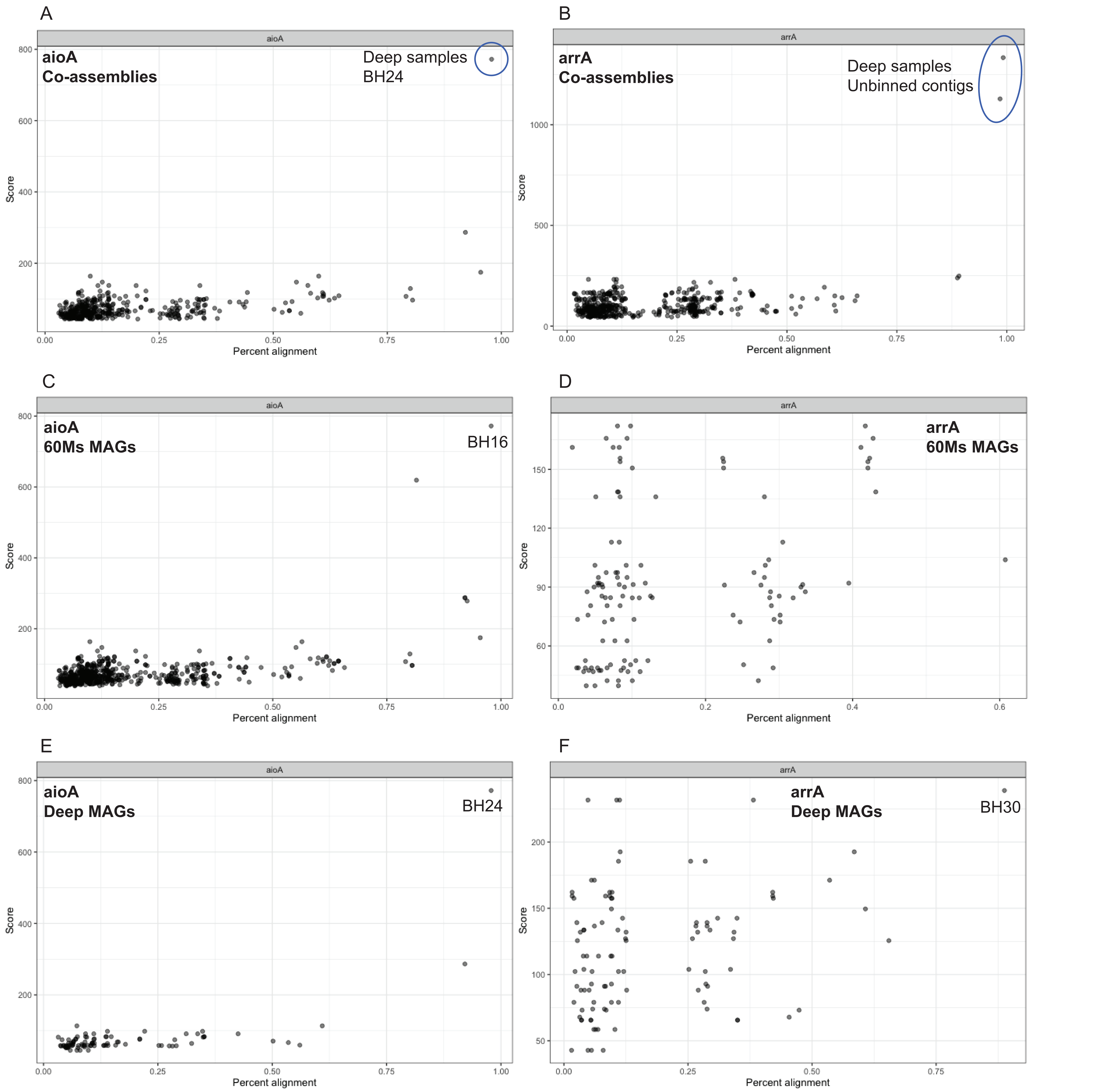
