## Supplemental Figure 3 for "Gulf of Mexico blue hole harbors high levels of novel microbial lineages"

Figure S3. Alpha diversity comparisons of depth groupings show similar Shannon diversity among all groups and higher Simpson diversity in the deepest group.

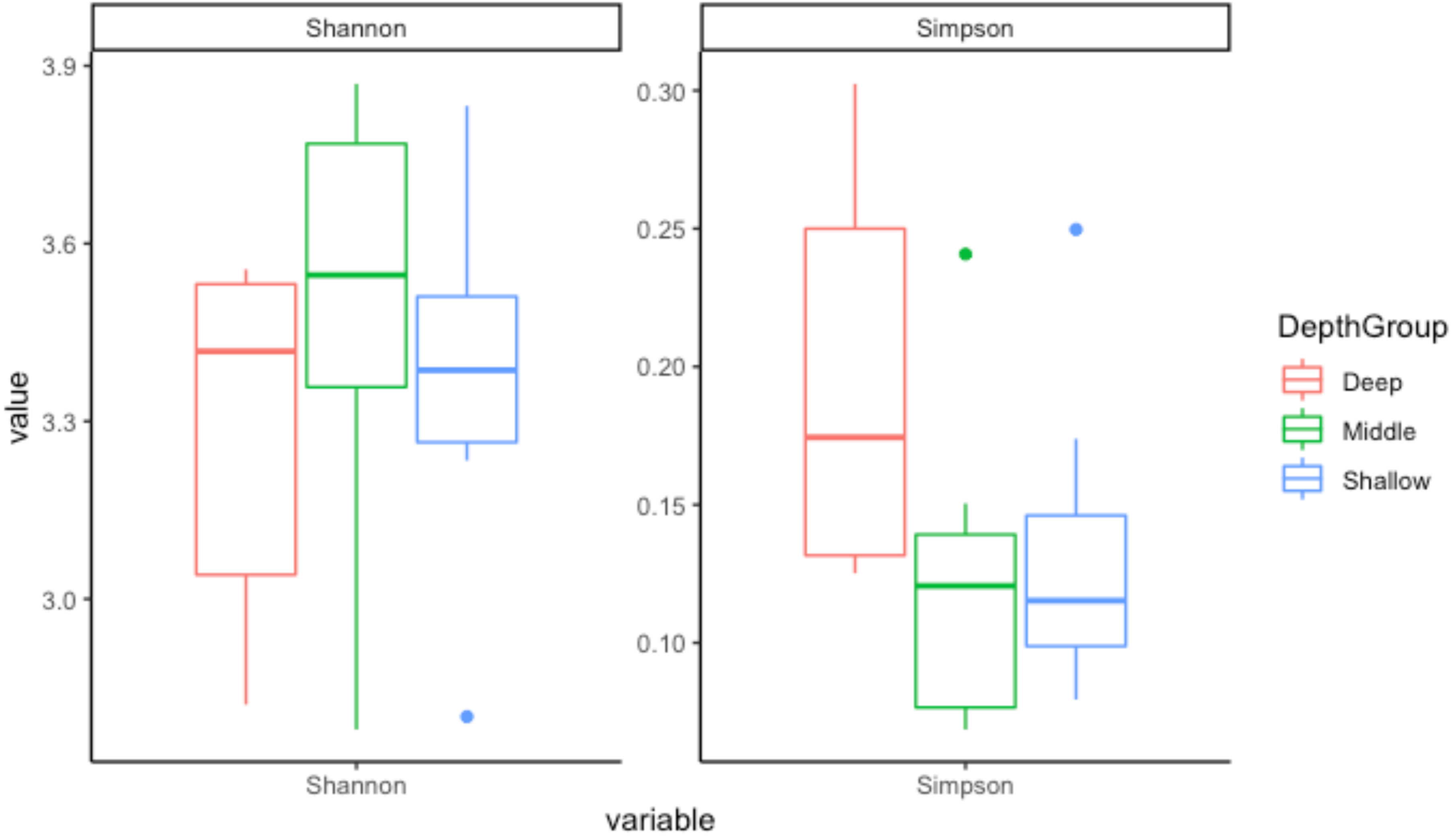
