## Supplemental Figure 7 for "Gulf of Mexico blue hole harbors high levels of novel microbial lineages"

Figure S7. Normalized abundance of two genes comprising the KEGG 'Signaling molecules and interaction' category in each pair of metagenomes.

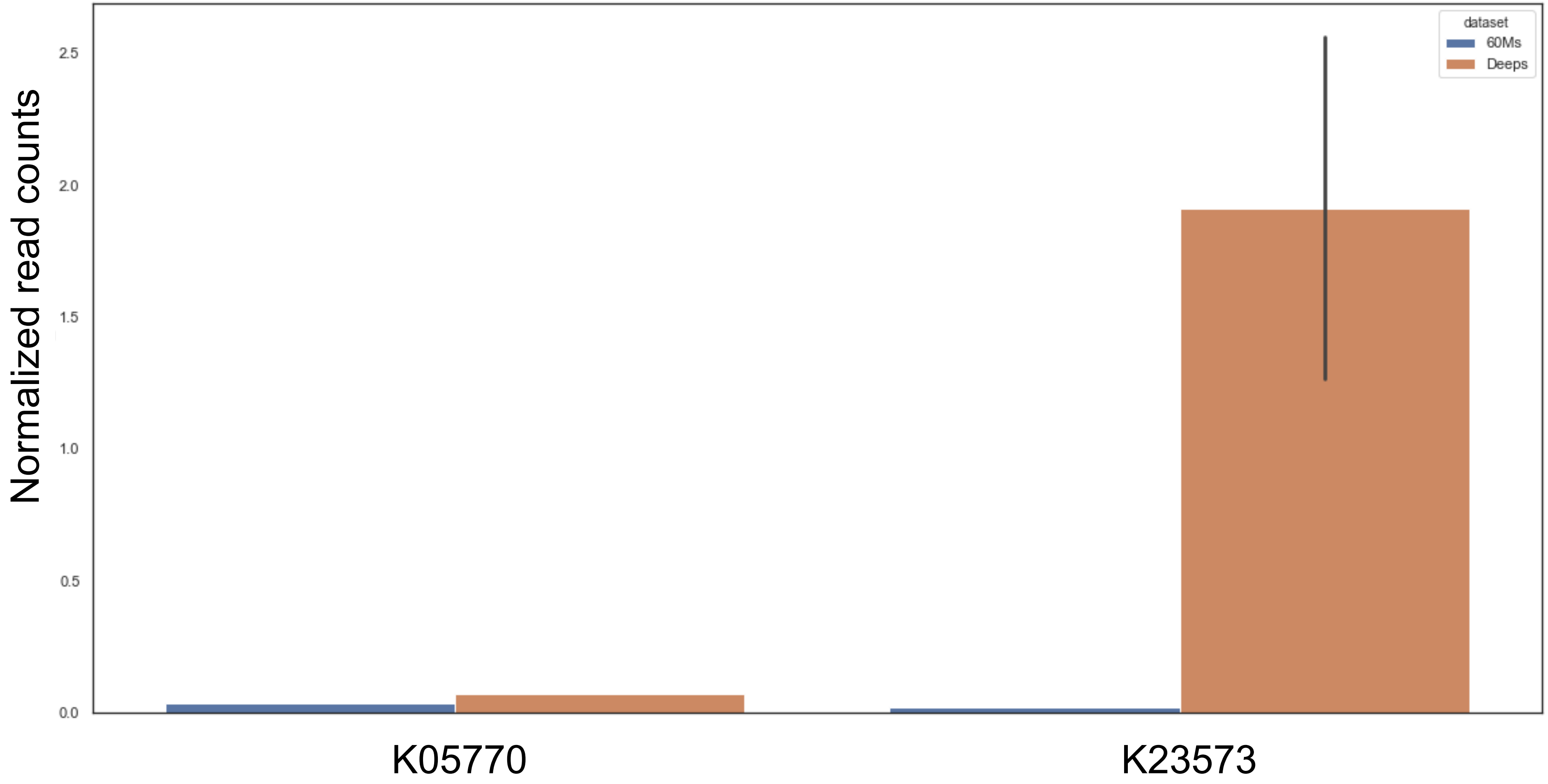
