## Supplemental Figure 9 for "Gulf of Mexico blue hole harbors high levels of novel microbial lineages"

Figure S8. Average cell counts at five depths sampled in September 2019. All counts represent duplicate filters except 50 m which had only one filter.

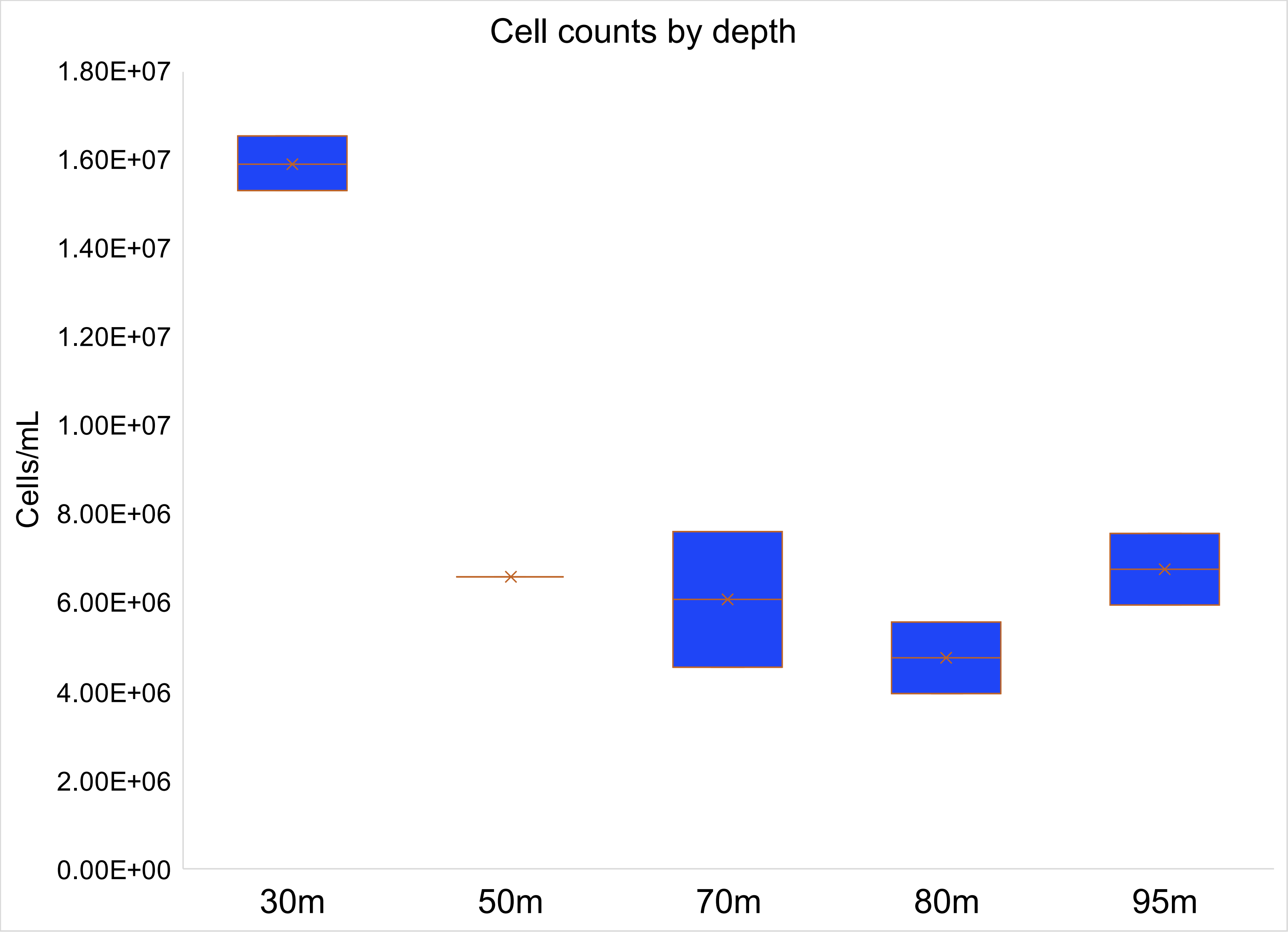
