## Supplemental Table 1 for "Gulf of Mexico blue hole harbors high levels of novel microbial lineages"

Table S1. Amberjack hole water column sample metadata and amplicon sequencing results. Processed reads are those that were quality-filtered, trimmed, and run through DADA2.

| Sample ID | Month | Depth (m) | Sampling method | # Raw reads | # processed reads |
|---|---|---|---|---|---|
| BH050M | May | 0 | Niskin (hand-cast) | 83,548 | 30,537 |
| BH057M | May | 7 | Niskin (rosette) | 84,483 | 40,768 |
| BH0515MA | May | 15 | Niskin (rosette) | 102,362 | 44,445 |
| BH0515MB | May | 15 | Diver bottle | 50,951 | 12,692 |
| BH0523M | May | 23 | Niskin (rosette) | 88,320 | 49,639 |
| BH0530M | May | 30 | Niskin (hand-cast) | 94,299 | 47,610 |
| BH0532C | May | 32 | Diver bottle | 79,530 | 39,941 |
| BH0546M | May | 46 | Diver bottle | 109,288 | 55,089 |
| BH0560M | May | 60 | Niskin (hand-cast) | 111,130 | 62,497 |
| BH0561M | May | 61 | Diver bottle | 75,200 | 33,385 |
| BH0585M | May | 85 | Niskin (hand-cast) | 116,106 | 59,356 |
| BH05106M | May | 106 | Diver bottle | 93,844 | 48,642 |
| BH0910M | September | 10 | Niskin (hand-cast) | 146,001 | 83,136 |
| BH0920M | September | 20 | Niskin (hand-cast) | 133,862 | 80,147 |
| BH0930M | September | 30 | Niskin (hand-cast) | 162,528 | 86,957 |
| BH0940M | September | 40 | Niskin (hand-cast) | 170,522 | 89,819 |
| BH0950M | September | 50 | Niskin (hand-cast) | 194,395 | 118,397 |
| BH0960M | September | 60 | Niskin (hand-cast) | 172,560 | 97,492 |
| BH0970M | September | 70 | Niskin (hand-cast) | 141,769 | 84,787 |
| BH0980M | September | 80 | Niskin (hand-cast) | 190,289 | 106,121 |
| BH0985M | September | 85 | Niskin (hand-cast) | 173,136 | 95,596 |
| BH0990M | September | 90 | Niskin (hand-cast) | 128,982 | 67,244 |
| BH0995M | September | 95 | Niskin (hand-cast) | 172,152 | 102,331 |
