## Supplemental Table 2 for "Gulf of Mexico blue hole harbors high levels of novel microbial lineages"

Table S2. Metagenome assembly data including sample source, pre- and post-quality filtered rea

| | | | | | SPAdes Ass | |
|---|---|---|---|---|---|---|
| **Sample ID** | **Month** | **Depth (m)** | **# Raw Reads** | **# Filtered Reads** | **# contigs > 1Kbp** | **N50** |
| BH0560M | May | 60 | 3,240,148 | 2,202,966 | 9,869 | 2,247 |
| BH05106M | May | 106 | 3,729,927 | 1,483,928 | 7,425 | 1,733 |
| BH0960M | September | 60 | 13,150,727 | 13,134,788 | 31,896 | 3,527 |
| BH0995M | September | 95 | 11,894,511 | 9,488,634 | 40,713 | 2,673 |

ads, and assembly statistics for both single and co-assemblies.

| embly | Co-assembly | | |
|---|---|---|---|
| longest contig (bp) | # contigs > 1Kbp | N50 | longest contig (bp) |
| 405,204 | 10,696 | 12,222 | 625,129 |
| 108,892 | 11,167 | 12,834 | 453,391 |
| 421,576 | 10,696 | 12,222 | 625,129 |
| 598,144 | 11,167 | 12,834 | 453,391 |
