## Supplemental Table 3 for "Gulf of Mexico blue hole harbors high levels of novel microbial lineages"

Table S3. Metagenome-assembled genomes (MAGs) and their associated samples,

| | | anvi'o quality metrics | | CheckM quality metrics | |
|---|---|---|---|---|---|
| **MAG ID** | **Depth (m)** | **Completeness** | **Redundancy** | **Completeness** | **Contamination** |
| BH1 | 60 | 94.51 | 0 | 95.77 | 0 |
| BH2 | 60 | 77.22 | 0.42 | 87.32 | 8.45 |
| BH3 | 60 | 82.4 | 4.24 | 86.84 | 5.26 |
| BH4 | 60 | 72.93 | 0.8 | 90.79 | 2.63 |
| BH5 | 60 | 98.25 | 7.18 | 100 | 15.49 |
| BH6 | 60 | 45.99 | 0 | 0 | 0 |
| BH7 | 60 | 97.44 | 8.12 | 92.96 | 5.63 |
| BH8 | 60 | 96.58 | 5.79 | 100 | 23.94 |
| BH9 | 60 | 96.55 | 9.91 | 97.18 | 23.94 |
| BH10 | 60 | 84.76 | 5.08 | 90.14 | 7.04 |
| BH11 | 60 | 80.4 | 7.14 | 90.79 | 2.63 |
| BH12 | 60 | 72.22 | 3.2 | 64.47 | 5.26 |
| BH13 | 60 | 86 | 1.6 | 93.42 | 1.32 |
| BH14 | 60 | 89.68 | 4.47 | 98.59 | 18.31 |
| BH15 | 60 | 82.36 | 0 | 96.05 | 1.32 |
| BH16 | 60 | 95.73 | 2.14 | 97.18 | 0 |
| BH17 | 60 | 73.42 | 2.58 | 67.61 | 4.23 |
| BH18 | 60 | 97.66 | 0.57 | 92.96 | 0 |
| BH19 | 60 | 53.12 | 6.45 | 63.16 | 5.26 |
| BH20 | 95-106 | 92.14 | 1.66 | 100 | 2.82 |
| BH21 | 95-106 | 73.36 | 0 | 67.11 | 1.32 |
| BH22 | 95-106 | 66.67 | 1.71 | 83.1 | 1.41 |
| BH23 | 95-106 | 94.57 | 0.54 | 91.55 | 1.41 |
| BH24 | 95-106 | 98.51 | 5.02 | 100 | 1.41 |
| BH25 | 95-106 | 96.77 | 1.61 | 98.59 | 2.82 |
| BH26 | 95-106 | 96.7 | 1.1 | 95.77 | 1.41 |
| BH27 | 95-106 | 92.24 | 1.9 | 77.46 | 2.82 |
| BH28 | 95-106 | 74.26 | 0.99 | 88.73 | 0 |
| BH29 | 95-106 | 95.1 | 3.86 | 87.32 | 9.86 |

| | | | | | |
|---|---|---|---|---|---|
| BH30 | 95-106 | 97.1 | 7.31 | 98.59 | 8.45 |
| BH31 | 95 | 53.15 | 1.4 | 81.69 | 5.63 |

quality metrics, and assigned taxonomy.

| Genome Taxonomy (GTDB) |
|---|
| d__Bacteria;p__Marinisomatota;c__Marinisomatia;o__Marinisomatales;f__TCS55;g__TCS55;s__ |
| d__Bacteria;p__Proteobacteria;c__Gammaproteobacteria;o__Pseudomonadales;f__Porticoccaceae;g__HTCC2207;s__ |
| d__Archaea;p__Thermoplasmatota;c__Poseidoniia;o__MGIII;f__CG-Epi1;g__CG-Epi1;s__ |
| d__Archaea;p__Thermoplasmatota;c__Poseidoniia;o__Poseidoniales;f__Thalassoarchaeaceae;g__MGIIb-O2;s__MGIIb-O2 sp002686525 |
| d__Bacteria;p__Proteobacteria;c__Gammaproteobacteria;o__Burkholderiales;f__Nitrosomonadaceae;g__GCA-2721545;s__ |
| d__Bacteria;p__Proteobacteria;c__Alphaproteobacteria;o__Puniceispirillales;f__Puniceispirillaceae;g__;s__ |
| d__Bacteria;p__Actinobacteriota;c__Acidimicrobiia;o__Microtrichales;f__TK06;g__UBA6944;s__UBA6944 sp002296525 |
| d__Bacteria;p__Actinobacteriota;c__Acidimicrobiia;o__Microtrichales;f__UBA11606;g__UBA11606;s__ |
| d__Bacteria;p__Proteobacteria;c__Gammaproteobacteria;o__Methylococcales;f__Methylomonadaceae;g__OPU3-GD-OMZ;s__ |
| d__Bacteria;p__Proteobacteria;c__Gammaproteobacteria;o__UBA10353;f__LS-SOB;g__UBA5682;s__ |
| d__Archaea;p__Thermoplasmatota;c__Poseidoniia;o__Poseidoniales;f__Thalassoarchaeaceae;g__MGIIb-O1;s__MGIIb-O1 sp002496905 |
| d__Archaea;p__Thermoplasmatota;c__Poseidoniia;o__Poseidoniales;f__Thalassoarchaeaceae;g__MGIIb-O1;s__MGIIb-O1 sp002502365 |
| d__Archaea;p__Thermoplasmatota;c__Poseidoniia;o__Poseidoniales;f__Thalassoarchaeaceae;g__MGIIb-O1;s__ |
| d__Bacteria;p__Proteobacteria;c__Gammaproteobacteria;o__Pseudomonadales;f__Nitrincolaceae;g__ASP10-02a;s__ |
| d__Archaea;p__Thermoplasmatota;c__Poseidoniia;o__Poseidoniales;f__Thalassoarchaeaceae;g__;s__ |
| d__Bacteria;p__Actinobacteriota;c__Acidimicrobiia;o__Microtrichales;f__MedAcidi-G1;g__UBA9410;s__ |
| d__Bacteria;p__Myxococcota;c__UBA796;o__UBA796;f__UBA796;g__;s__ |
| d__Bacteria;p__Planctomycetota;c__Phycisphaerae;o__Phycisphaerales;f__SM1A02;g__GCA-002718515;s__ |
| d__Archaea;p__Crenarchaeota;c__Nitrososphaeria;o__Nitrososphaerales;f__Nitrosopumilaceae;g__Nitrosopumilus;s__ |
| d__Bacteria;p__Proteobacteria;c__Gammaproteobacteria;o__Thiomicrospirales;f__Thioglobaceae;g__UBA2013;s__ |
| d__Archaea;p__Nanoarchaeota;c__Nanoarchaeia;o__SCGC-AAA011-G17;f__UBA489;g__;s__ |
| d__Bacteria;p__Patescibacteria;c__ABY1;o__SG8-24;f__UBA11717;g__UBA11717;s__ |
| d__Bacteria;p__Bacteroidota;c__Bacteroidia;o__Bacteroidales;f__F082;g__;s__ |
| d__Bacteria;p__Proteobacteria;c__Alphaproteobacteria;o__Rhodospirillales_A;f__UBA3470;g__TMED8;s__ |
| d__Bacteria;p__Bacteroidota;c__Bacteroidia;o__Bacteroidales;f__F082;g__GCA-002708315;s__ |
| d__Bacteria;p__Marinisomatota;c__UBA8477;o__UBA8477;f__UBA8477;g__;s__ |
| d__Bacteria;p__Marinisomatota;c__UBA8477;o__UBA8477;f__UBA8477;g__;s__ |
| d__Bacteria;p__Patescibacteria;c__ABY1;o__SG8-24;f__GWF2-40-263;g__UM-FILTER-50-9;s__ |
| d__Bacteria;p__Marinisomatota;c__UBA8477;o__UBA8477;f__UBA8477;g__;s__ |

d__Bacteria;p__Desulfobacterota;c__Desulfobacteria;o__Desulfatiglandales;f__NaphS2;g__NaphS2;s__
d__Bacteria;p__Myxococcota;c__UBA796;o__UBA796;f__;g__;s__

| 16S rRN |
| --- |
| Marinimicrobia (SAR406 clade); D_2__uncultured bacterium HF0500_01L02; D_3__uncultured bacterium HF0500_01L02; D_4__uncultured bacterium HF050 |
| Proteobacteria; D_2__Gammaproteobacteria; D_3__Cellvibrionales; D_4__Porticoccaceae; D_5__SAR92 clade |
| Euryarchaeota; D_2__Thermoplasmata; D_3__Marine Group II; D_4__uncultured marine archaeon DCM3921; D_5__uncultured marine archaeon DCM3921; |
| Proteobacteria; D_2__Gammaproteobacteria; D_3__Betaproteobacteriales; D_4__Nitrosomonadaceae; D_5__Nitrosomonas; D_6__marine metagenome |
| Proteobacteria; D_2__Gammaproteobacteria; D_3__SAR86 clade |
| Euryarchaeota; D_2__Thermoplasmata; D_3__Marine Group II; D_4__uncultured marine group II/III euryarchaeote KM3_53_G07; D_5__uncultured marine gr |
| Planctomycetes; D_2__Phycisphaerae; D_3__Phycisphaerales; D_4__Phycisphaeraceae; D_5__JL-ETNP-F27 |
| Thaumarchaeota; D_2__Nitrososphaeria; D_3__Nitrosopumilales; D_4__Nitrosopumilaceae; D_5__Candidatus Nitrosopumilus |
| Proteobacteria; D_2__Gammaproteobacteria; D_3__Thiomicrospirales; D_4__Thioglobaceae; D_5__SUP05 cluster; D_6__uncultured gamma proteobacteriur |
| Nanoarchaeaeota; D_2__Woesearchaeia |
| (1) Patescibacteria; D_2__ABY1; D_3__Candidatus Uhrbacteria; (2) D_0__Bacteria; D_1__Planctomycetes; D_2__Phycisphaerae |
| Proteobacteria; D_2__Alphaproteobacteria; D_3__Rhodospirillales; D_4__Magnetospiraceae; D_5__uncultured; D_6__uncultured alpha proteobacterium |
| Bacteroidetes;D_2__Bacteroidia;D_3__Bacteroidetes VC2.1 Bac22;D_4__uncultured alpha proteobacterium;D_5__uncultured alpha proteobacterium;D_6__ui |
