## Supplemental Table 4 for "Gulf of Mexico blue hole harbors high levels of novel microbial lineages"

Table S4. MAGs used for the phylogenomic trees (Fig. 4, Fig. S4, Fig. S5) including the blue hole MAG of interest and the source of all other MAGs included in the phylogeny.

| Phylogeny | | | | |
|---|---|---|---|---|
| Figure | Blue Hole MAG | Database | Accession #/Taxon ID | Source |
| 4 | Nitrosopumilus sp. (BH19) | FigShare (Aylward Lab) | HMT_AABW | marine water (>4K m depth) |
| 4 | Nitrosopumilus sp. (BH19) | FigShare (Aylward Lab) | HMT_AAIW | marine water (>4K m depth) |
| 4 | Nitrosopumilus sp. (BH19) | FigShare (Aylward Lab) | HMT_ATL | marine water (>4K m depth) |
| 4 | Nitrosopumilus sp. (BH19) | FigShare (Aylward Lab) | HMT_NADW | marine water (>4K m depth) |
| 4 | Nitrosopumilus sp. (BH19) | FigShare (Aylward Lab) | HMT_PAC | marine water (>4K m depth) |
| 4 | Nitrosopumilus sp. (BH19) | GenBank | GCA_011058825.1 | hot spring sediment |
| 4 | Nitrosopumilus sp. (BH19) | GenBank | GCA_002906215.1 | hot spring biofilm |
| 4 | Nitrosopumilus sp. (BH19) | GenBank | GCA_900248165.1 | terrestrial hot spring |
| 4 | Nitrosopumilus sp. (BH19) | GenBank | GCA_001627015.1 | Red Sea water column |
| 4 | Nitrosopumilus sp. (BH19) | GenBank | GCA_001627075.1 | Red Sea water column |
| 4 | Nitrosopumilus sp. (BH19) | GenBank | GCA_001627095.1 | Red Sea water column |
| 4 | Nitrosopumilus sp. (BH19) | GenBank | GCA_001627105.1 | Red Sea water column |
| 4 | Nitrosopumilus sp. (BH19) | GenBank | GCA_001627165.1 | Red Sea water column |
| 4 | Nitrosopumilus sp. (BH19) | GenBank | GCA_001627235.1 | Red Sea water column |
| 4 | Nitrosopumilus sp. (BH19) | GenBank | GCA_001437625.1 | brackish water |
| 4 | Nitrosopumilus sp. (BH19) | GenBank | GCA_001438895.1 | brackish water |
| 4 | Nitrosopumilus sp. (BH19) | GenBank | GCA_001541925.1 | cold seep sponge |
| 4 | Nitrosopumilus sp. (BH19) | GenBank | GCA_001543015.1 | glass sponge |
| 4 | Nitrosopumilus sp. (BH19) | GenBank | GCA_001674955.1 | marine water |
| 4 | Nitrosopumilus sp. (BH19) | GenBank | GCA_000018465 | isolate |
| 4 | Nitrosopumilus sp. (BH19) | GenBank | GCA_000242875 | coastal sediments |
| 4 | Nitrosopumilus sp. (BH19) | GenBank | GCA_002494985 | deep-sea sponge |
| 4 | Nitrosopumilus sp. (BH19) | GenBank | GCA_002506665 | deep-sea sponge |
| 4 | Nitrosopumilus sp. (BH19) | GenBank | GCA_002508395 | water |
| 4 | Nitrosopumilus sp. (BH19) | GenBank | GCA_002690535 | marine water |
| 4 | Nitrosopumilus sp. (BH19) | GenBank | GCA_002763195 | groundwater |
| 4 | Nitrosopumilus sp. (BH19) | GenBank | GCA_000299365 | isolate |
| 4 | Nitrosopumilus sp. (BH19) | GenBank | GCA_000299395 | isolate |
| 4 | Nitrosopumilus sp. (BH19) | GenBank | GCA_003175215 | isolate |

| | | | | |
|---|---|---|---|---|
| 4 | Nitrosopumilus sp. (BH19) | GenBank | GCA_003331425 | marine water |
| 4 | Nitrosopumilus sp. (BH19) | GenBank | GCA_003702495 | marine water |
| 4 | Nitrosopumilus sp. (BH19) | GenBank | GCA_003702525 | marine water |
| 4 | Nitrosopumilus sp. (BH19) | GenBank | GCA_003702545 | marine water |
| 4 | Nitrosopumilus sp. (BH19) | GenBank | GCA_003724235 | isolate |
| 4 | Nitrosopumilus sp. (BH19) | GenBank | GCA_003724255 | isolate |
| 4 | Nitrosopumilus sp. (BH19) | GenBank | GCA_003724285 | isolate |
| 4 | Nitrosopumilus sp. (BH19) | GenBank | GCA_003724325 | isolate |
| 4 | Nitrosopumilus sp. (BH19) | GenBank | GCA_006740685 | marine water |
| 4 | Nitrosopumilus sp. (BH19) | GenBank | GCA_007571005 | PNG CO2 seep-seep site |
| 4 | Nitrosopumilus sp. (BH19) | GenBank | GCA_007571135 | PNG CO2 seep-control site |
| 4 | Nitrosopumilus sp. (BH19) | GenBank | GCA_008080815 | marine water |
| 4 | Nitrosopumilus sp. (BH19) | GenBank | GCA_008080855 | marine water |
| 4 | Nitrosopumilus sp. (BH19) | GenBank | GCA_000875775 | surface seawater |
| 4 | Nitrosopumilus sp. (BH19) | GenBank | GCA_000956175 | marine water |
| 4 | Nitrosopumilus sp. (BH19) | GenBank | GCA_009694705 | freshwater |
| 4 | Nitrosopumilus sp. (BH19) | FigShare (Patin) | Nitrosopumilus_manual | Amberjack hole (this study) |
| 4 | Nitrosopumilus sp. (BH19) | GenBank | GCA_000328945.1 | marine sediment |
| 4 | Nitrosopumilus sp. (BH19) | GenBank | GCA_000698785.1 | isolate |
| 4 | Nitrosopumilus sp. (BH19) | GenBank | GCA_000189555.1 | isolate |
| S4 | SUP05 (Thioglobaceae) (BH20) | GenBank | GCA_001281385.1 | Puget Sound water |
| S4 | SUP05 (Thioglobaceae) (BH20) | GenBank | GCA_001293165.1 | Effingham Inlet water |
| S4 | SUP05 (Thioglobaceae) (BH20) | GenBank | GCA_001628345.1 | Red Sea water |
| S4 | SUP05 (Thioglobaceae) (BH20) | GenBank | GCA_001628365.1 | Red Sea water |
| S4 | SUP05 (Thioglobaceae) (BH20) | GenBank | GCA_001628405.1 | Red Sea water |
| S4 | SUP05 (Thioglobaceae) (BH20) | GenBank | GCA_001682155.1 | Puget Sound water |
| S4 | SUP05 (Thioglobaceae) (BH20) | GenBank | GCA_000205985.2 | Saanich Inlet |
| S4 | SUP05 (Thioglobaceae) (BH20) | GenBank | GCA_002169705.1 | marine water |
| S4 | SUP05 (Thioglobaceae) (BH20) | GenBank | GCA_002456985.1 | marine water |
| S4 | SUP05 (Thioglobaceae) (BH20) | GenBank | GCA_002456995.1 | marine water |
| S4 | SUP05 (Thioglobaceae) (BH20) | GenBank | GCA_002689885.1 | marine water |
| S4 | SUP05 (Thioglobaceae) (BH20) | GenBank | GCA_002909145.1 | Peruvian upwelling |
| S4 | SUP05 (Thioglobaceae) (BH20) | GenBank | GCA_003326015.1 | marine water |

| | | | | |
|---|---|---|---|---|
| S4 | SUP05 (Thioglobaceae) (BH20) | GenBank | GCA_003332345.1 | marine water |
| S4 | SUP05 (Thioglobaceae) (BH20) | GenBank | GCA_003978155.1 | hydrothermal vent |
| S4 | SUP05 (Thioglobaceae) (BH20) | GenBank | GCA_003978165.1 | hydrothermal vent |
| S4 | SUP05 (Thioglobaceae) (BH20) | GenBank | GCA_003978175.1 | hydrothermal vent |
| S4 | SUP05 (Thioglobaceae) (BH20) | GenBank | GCA_003978185.1 | hydrothermal vent |
| S4 | SUP05 (Thioglobaceae) (BH20) | GenBank | GCA_003978205.1 | hydrothermal vent |
| S4 | SUP05 (Thioglobaceae) (BH20) | GenBank | GCA_003978255.1 | hydrothermal vent |
| S4 | SUP05 (Thioglobaceae) (BH20) | GenBank | GCA_003978285.1 | hydrothermal vent |
| S4 | SUP05 (Thioglobaceae) (BH20) | GenBank | GCA_003978295.1 | hydrothermal vent |
| S4 | SUP05 (Thioglobaceae) (BH20) | GenBank | GCA_004212795.1 | marine water |
| S4 | SUP05 (Thioglobaceae) (BH20) | GenBank | GCA_004212825.1 | marine water |
| S4 | SUP05 (Thioglobaceae) (BH20) | GenBank | GCA_000424685.1 | isolate |
| S4 | SUP05 (Thioglobaceae) (BH20) | GenBank | GCA_008364125.1 | hydrothermal vent sponge |
| S4 | SUP05 (Thioglobaceae) (BH20) | GenBank | GCA_008364135.1 | hydrothermal vent sponge |
| S4 | SUP05 (Thioglobaceae) (BH20) | FigShare (Patin) | BH_Deeps_001 | Amberjack hole (this study) |
| S4 | SUP05 (Thioglobaceae) (BH20) | GenBank | GCA_900180385.1 | gill tissue |
| S4 | SUP05 (Thioglobaceae) (BH20) | GenBank | GCA_900180335.1 | gill tissue |
| S4 | SUP05 (Thioglobaceae) (BH20) | GenBank | GCA_900180375.1 | sponge tissue |
| S4 | SUP05 (Thioglobaceae) (BH20) | GenBank | GCA_900180405.1 | sponge tissue |
| S4 | SUP05 (Thioglobaceae) (BH20) | GenBank | GCA_900180365.1 | sponge tissue |
| S5 | Woesearchaeota (BH21) | GenBank | GCA_000018465 | isolate |
| S5 | Woesearchaeota (BH21) | GenBank | GCA_902529885.1 | hot spring sediment |
| S5 | Woesearchaeota (BH21) | GenBank | GCA_011045925.1 | hot spring sediment |
| S5 | Woesearchaeota (BH21) | GenBank | GCA_011331795.1 | mud sediment |
| S5 | Woesearchaeota (BH21) | GenBank | GCA_011333755.1 | mud sediment |
| S5 | Woesearchaeota (BH21) | GenBank | GCA_011333955.1 | hot spring sediment |
| S5 | Woesearchaeota (BH21) | GenBank | GCA_011368255.1 | deep-sea hydrothermal vent sediment |
| S5 | Woesearchaeota (BH21) | GenBank | GCA_011372335.1 | mud sediment |
| S5 | Woesearchaeota (BH21) | GenBank | GCA_011374165.1 | contaminated sediment |
| S5 | Woesearchaeota (BH21) | GenBank | GCA_001791825.1 | groundwater |
| S5 | Woesearchaeota (BH21) | GenBank | GCA_001871415.1 | groundwater |
| S5 | Woesearchaeota (BH21) | GenBank | GCA_001872765.1 | groundwater |
| S5 | Woesearchaeota (BH21) | GenBank | GCA_001872825.1 | soil |

| | | | | |
|---|---|---|---|---|
| S5 | Woesearchaeota (BH21) | GenBank | GCA_002498125.1 | water |
| S5 | Woesearchaeota (BH21) | GenBank | GCA_002503425.1 | waste water |
| S5 | Woesearchaeota (BH21) | GenBank | GCA_002503705.1 | marine water |
| S5 | Woesearchaeota (BH21) | GenBank | GCA_002503745.1 | marine OMZ (Arabian Sea) |
| S5 | Woesearchaeota (BH21) | GenBank | GCA_002505585.1 | marine water |
| S5 | Woesearchaeota (BH21) | GenBank | GCA_002505845.1 | crude oil |
| S5 | Woesearchaeota (BH21) | GenBank | GCA_002505945.1 | marine OMZ (Arabian Sea) |
| S5 | Woesearchaeota (BH21) | GenBank | GCA_002506165.1 | marine water |
| S5 | Woesearchaeota (BH21) | GenBank | GCA_002685855.1 | marine water |
| S5 | Woesearchaeota (BH21) | GenBank | GCA_002686215.1 | marine water |
| S5 | Woesearchaeota (BH21) | GenBank | GCA_002686295.1 | marine water |
| S5 | Woesearchaeota (BH21) | GenBank | GCA_002686855.1 | marine water |
| S5 | Woesearchaeota (BH21) | GenBank | GCA_002687275.1 | marine water |
| S5 | Woesearchaeota (BH21) | GenBank | GCA_002687795.1 | marine water |
| S5 | Woesearchaeota (BH21) | GenBank | GCA_002688315.1 | marine water |
| S5 | Woesearchaeota (BH21) | GenBank | GCA_002688775.1 | marine water |
| S5 | Woesearchaeota (BH21) | GenBank | GCA_002690055.1 | marine water |
| S5 | Woesearchaeota (BH21) | GenBank | GCA_002725495.1 | groundwater |
| S5 | Woesearchaeota (BH21) | GenBank | GCA_002762695.1 | groundwater |
| S5 | Woesearchaeota (BH21) | GenBank | GCA_002762705.1 | groundwater |
| S5 | Woesearchaeota (BH21) | GenBank | GCA_002762765.1 | groundwater |
| S5 | Woesearchaeota (BH21) | GenBank | GCA_002762785.1 | groundwater |
| S5 | Woesearchaeota (BH21) | GenBank | GCA_002762795.1 | groundwater |
| S5 | Woesearchaeota (BH21) | GenBank | GCA_002762845.1 | groundwater |
| S5 | Woesearchaeota (BH21) | GenBank | GCA_002762855.1 | groundwater |
| S5 | Woesearchaeota (BH21) | GenBank | GCA_002762865.1 | groundwater |
| S5 | Woesearchaeota (BH21) | GenBank | GCA_002762915.1 | groundwater |
| S5 | Woesearchaeota (BH21) | GenBank | GCA_002762925.1 | groundwater |
| S5 | Woesearchaeota (BH21) | GenBank | GCA_002762985.1 | groundwater |
| S5 | Woesearchaeota (BH21) | GenBank | GCA_002763025.1 | groundwater |
| S5 | Woesearchaeota (BH21) | GenBank | GCA_002763335.1 | groundwater |
| S5 | Woesearchaeota (BH21) | GenBank | GCA_002779235.1 | groundwater |
| S5 | Woesearchaeota (BH21) | GenBank | GCA_002780105.1 | groundwater |

| | | | | |
|---|---|---|---|---|
| S5 | Woesearchaeota (BH21) | GenBank | GCA_002784405.1 | groundwater |
| S5 | Woesearchaeota (BH21) | GenBank | GCA_002792055.1 | groundwater |
| S5 | Woesearchaeota (BH21) | GenBank | GCA_002792075.1 | groundwater |
| S5 | Woesearchaeota (BH21) | GenBank | GCA_002792115.1 | groundwater |
| S5 | Woesearchaeota (BH21) | GenBank | GCA_002794135.1 | enrichment from estuary sediment |
| S5 | Woesearchaeota (BH21) | GenBank | GCA_002867475.1 | hypersaline soda lake sediment |
| S5 | Woesearchaeota (BH21) | GenBank | GCA_003555225.1 | hypersaline soda lake sediment |
| S5 | Woesearchaeota (BH21) | GenBank | GCA_003558875.1 | hypersaline soda lake sediment |
| S5 | Woesearchaeota (BH21) | GenBank | GCA_003560545.1 | hypersaline soda lake sediment |
| S5 | Woesearchaeota (BH21) | GenBank | GCA_003561825.1 | hypersaline soda lake sediment |
| S5 | Woesearchaeota (BH21) | GenBank | GCA_003564925.1 | subsurface aquifer |
| S5 | Woesearchaeota (BH21) | GenBank | GCA_003599055.1 | subsurface aquifer |
| S5 | Woesearchaeota (BH21) | GenBank | GCA_003599145.1 | deep-sea hydrothermal vent sediment |
| S5 | Woesearchaeota (BH21) | GenBank | GCA_003649025.1 | deep-sea hydrothermal vent sediment |
| S5 | Woesearchaeota (BH21) | GenBank | GCA_003649035.1 | deep-sea hydrothermal vent sediment |
| S5 | Woesearchaeota (BH21) | GenBank | GCA_003650545.1 | deep-sea hydrothermal vent sediment |
| S5 | Woesearchaeota (BH21) | GenBank | GCA_003650585.1 | iron-rich hot spring |
| S5 | Woesearchaeota (BH21) | GenBank | GCA_003694385.1 | iron-rich hot spring |
| S5 | Woesearchaeota (BH21) | GenBank | GCA_003694495.1 | iron-rich hot spring |
| S5 | Woesearchaeota (BH21) | GenBank | GCA_003694805.1 | iron-rich hot spring |
| S5 | Woesearchaeota (BH21) | GenBank | GCA_003695045.1 | iron-rich hot spring |
| S5 | Woesearchaeota (BH21) | GenBank | GCA_003695265.1 | iron-rich hot spring |
| S5 | Woesearchaeota (BH21) | GenBank | GCA_003695435.1 | marine sediment |
| S5 | Woesearchaeota (BH21) | GenBank | GCA_005222965.1 | hypersaline soda lake brine |
| S5 | Woesearchaeota (BH21) | GenBank | GCA_007116295.1 | hypersaline soda lake brine |
| S5 | Woesearchaeota (BH21) | GenBank | GCA_007116315.1 | hypersaline soda lake brine |
| S5 | Woesearchaeota (BH21) | GenBank | GCA_007116645.1 | hypersaline soda lake brine |
| S5 | Woesearchaeota (BH21) | GenBank | GCA_007116655.1 | hypersaline soda lake brine |
| S5 | Woesearchaeota (BH21) | GenBank | GCA_007116795.1 | hypersaline soda lake brine |
| S5 | Woesearchaeota (BH21) | GenBank | GCA_007117065.1 | hypersaline soda lake brine |
| S5 | Woesearchaeota (BH21) | GenBank | GCA_007117145.1 | hypersaline soda lake brine |
| S5 | Woesearchaeota (BH21) | GenBank | GCA_007117405.1 | hypersaline soda lake brine |
| S5 | Woesearchaeota (BH21) | GenBank | GCA_007117735.1 | hypersaline soda lake sediment |

| | | | | |
|---|---|---|---|---|
| S5 | Woesearchaeota (BH21) | GenBank | GCA_007128245.1 | hypersaline soda lake sediment |
| S5 | Woesearchaeota (BH21) | GenBank | GCA_007130955.1 | hypersaline soda lake sediment |
| S5 | Woesearchaeota (BH21) | GenBank | GCA_007131205.1 | hypersaline soda lake sediment |
| S5 | Woesearchaeota (BH21) | GenBank | GCA_007133845.1 | hypersaline soda lake sediment |
| S5 | Woesearchaeota (BH21) | GenBank | GCA_009691355.1 | freshwater |
| S5 | Woesearchaeota (BH21) | GenBank | GCA_902385765.1 | groundwater |
| S5 | Woesearchaeota (BH21) | IMG | 2806311055 | Black Sea water |
| S5 | Woesearchaeota (BH21) | IMG | 2806311056 | Black Sea water |
| S5 | Woesearchaeota (BH21) | IMG | 2785511131 | Black Sea water |
| S5 | Woesearchaeota (BH21) | IMG | 2785511134 | Black Sea water |
| S5 | Woesearchaeota (BH21) | IMG | 2811994887 | Black Sea water |
| S5 | Woesearchaeota (BH21) | FigShare (Patin) | BH_Deeps_002 | Amberjack hole (this study) |

Outgroup?

X

X

X
